# Collective cancer cell calcium activity drives brain metastasis

**DOI:** 10.64898/2026.05.10.723715

**Authors:** Nils R. Hebach, Niels Olshausen, Joana Schlag, Chanté D. Mayer, Jonas Henkenjohann, Theresa Kraft, Severin Bang, Calvin Thommek, Christina Nürnberg, Nico E. Horsak, Dirk C. Hoffmann, Alexandros Kourtesakis, Nicola Porzberg, Andres Flores Valle, C. Zoë Linke, Yvonne Yang, Daniel D. Azorín, David Hausmann, Varun Venkataramani, Friedegund Meier, Miriam Ratliff, Michael Seifert, Dana Westphal, Michael O. Breckwoldt, Jens Timmer, Kai Johnsson, Wolfgang Wick, Frank Winkler, Matthia A. Karreman

## Abstract

Communication in multicellular networks is a cancer-intrinsic neural feature and crucial for primary brain tumor growth and resistance, but it is unclear whether brain metastases (BrM), the most common and deadliest brain malignancy, are also driven by communicating cancer networks. Using intravital two-photon microscopy in awake mice, clinical specimens, and Ca^2+^ integrators, we demonstrate that brain-colonizing breast and lung cancer and melanoma cells display gap-junction-dependent, coordinated Ca^2+^ activity in multicellular, cancer-cell intrinsic networks, which drives their proliferation. Mechanistically, Ca^2+^ oscillations induce transcription of immediate early genes, adoption of a neuronal expression profile, and cell cycle progression. While many of those features are enriched in BrM, all investigated cancer cell lines showed collective Ca^2+^ activity. Therapeutically, blocking Ca^2+^ activity with gap junction inhibitors reduces BrM burden in mouse models. Here we show communicating cancer cell syncytia as drivers of BrM growth, pointing to a targetable pathomechanism, and potentially a new pan-cancer hallmark.

**Graphical Abstract:** In brief

Brain metastases form gap-junction-coupled networks exhibiting spontaneous, coordinated Ca^2+^ activity linked to immediate early gene activation, neuronal gene programs, and cell cycle progression. Disrupting Ca^2+^ network communication with gap junction inhibitors induces cell cycle arrest and reduces brain metastatic burden *in vivo*.

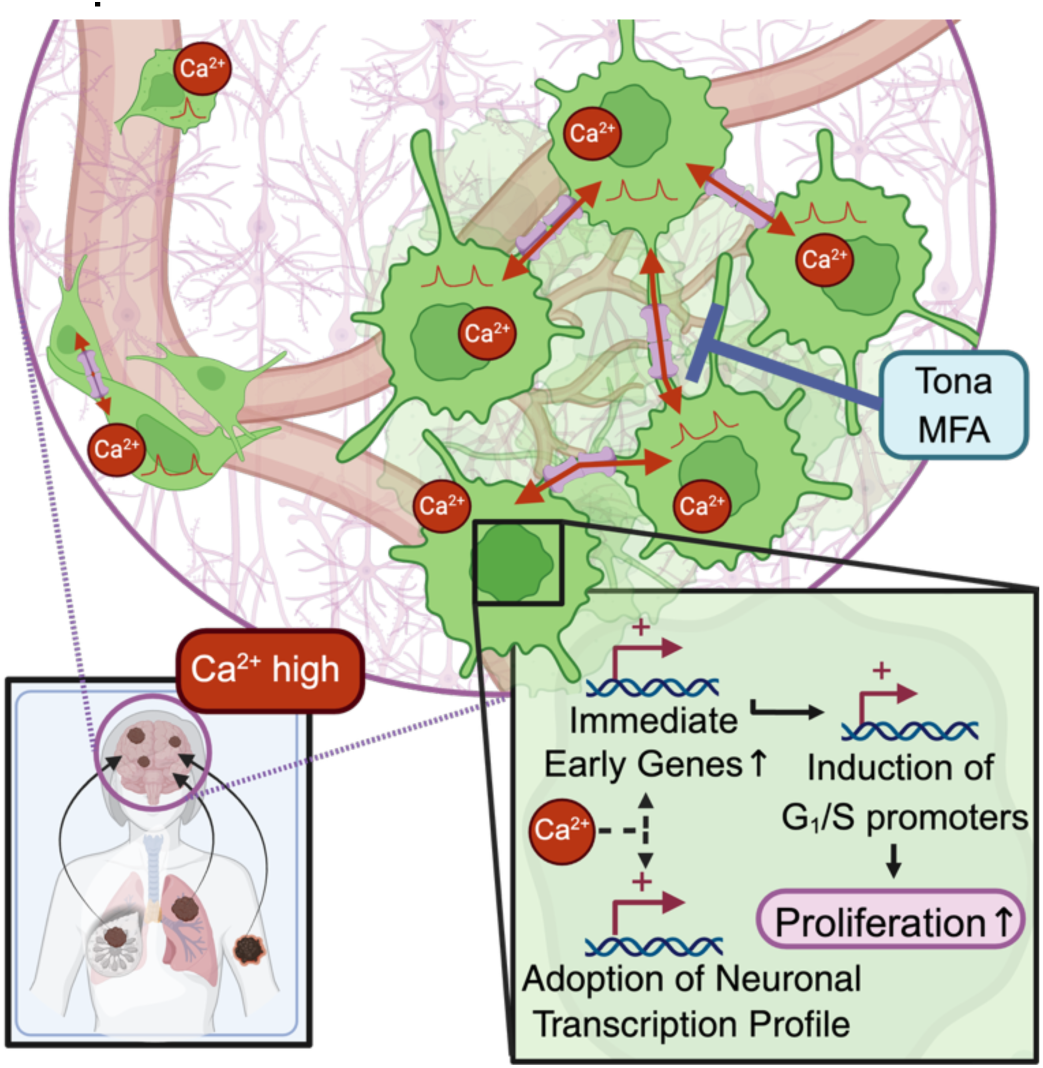

**Highlights:**

- Brain metastases display collective Ca^2+^ activity in gap-junction-coupled networks
- Ca^2+^ co-activity is conserved across cancers but enriched in brain metastasis
- Ca^2+^ oscillations robustly induce neuronal gene programs and cell cycle progression
- Gap junction inhibition reduces brain metastasis burden in mice

## Introduction

Research in the rapidly evolving field of Cancer Neuroscience has established that the nervous system actively drives tumor progression, metastasis, and therapeutic resistance through paracrine signaling^1–6^ and direct electrochemical communication^7–12^. Beyond responding to neural signals, cancer cells can themselves acquire neuronal or at least neural features^13,14^. This phenotypic mimicry has been well established in brain tumors, where a portion of tumor cells recapitulate transcriptional profiles of neural progenitor cells^10,15,16^.

Similarly, extracranial tumors can adopt a neuroendocrine differentiation profile^17–20^. Moreover, expression of neural gene signatures, such as axon-guidance programs^21,22^ and neurotransmitter signaling^23,24^, are associated with highly aggressive behavior such as metastasis, immunosuppression and therapy resistance^25^. Notably, brain metastases (BrM) from extracranial primary tumors, including melanoma and lung cancer, acquire neuronal-like transcriptional profiles upon brain colonization^26–30^ and can engage in gap-junction mediated crosstalk with astrocytes^31^. Together, this suggests that a certain level of neuronal identity of cancer cells of non-neural cancer entities may be either selected for during brain colonization - or actively induced by the brain microenvironment^32^.

Whether this neuro-cancer cell plasticity reprogramming potentially endows BrM with functional neuronal-like properties remains an open question. In primary brain tumors, such properties have well-documented consequences: glioma cells hijack neuronal mechanisms of invasion^10^ and upregulate axon-associated molecules to extend long membrane protrusions^33–35^. This enables the formation of functionally connected, malignant cell networks^12,35–37^ that coordinate the exchange of calcium ions (Ca^2+^) across the tumor mass, promoting progression and resistance to therapy^35,38^. These oscillations are sustained by a pacemaker-like subpopulation defined by expression of the Ca^2+^-activated potassium channel KCa3.1^39^. Similarly, neural cells use coordinated Ca^2+^ signaling for population-level network organization and memory^40,41^, and intercellular Ca^2+^signalling regulates gene-expression of *FOS* and other immediate early genes (IEG)^42,43^, which in turn influence circuit strength and neuronal plasticity. BrM are known to receive Ca^2+^ influx via synaptic^7,8,11^ and peri-synaptic^44^ communication with neurons. However, whether metastatic cells from carcinomas and melanomas also utilize cancer cell-intrinsic Ca^2+^ signaling, including the capacity to form organized, communicating networks, has not been investigated. Addressing this gap is particularly important given the dismal prognosis of BrM^45^ and the lack of long-term effective, targeted therapies in patients.

Here, we demonstrate that brain-metastatic cancer cells form neural-like, cancer-cell intrinsic gap junction-coupled networks exhibiting spontaneous, synchronized Ca^2+^ activity. Functional and molecular analyses identify Ca^2+^ signaling as a regulator of proliferation through cell cycle control, operating via transcription of immediate early genes and cell cycle progression. While coordinated Ca^2+^ activity and its transcriptional consequences are enriched in BrM relative to extracranial tumors, Ca^2+^ activity is detectable across all cancer entities examined, suggesting a broader role in cancer biology. Pharmacological disruption of network communication attenuates tumor growth, pointing to Ca^2+^-mediated intercellular signaling as a potential therapeutic vulnerability in BrM.

## Results

### Brain Metastatic Cancer Cells Display Spontaneous Ca^2+^ Activity

To gain insights into the occurrence and role of Ca^2+^ transients during the development of BrM *in vivo*, we performed intravital microscopy in awake mice following intracardiac injection of genetically encoded Ca^2+^ indicator-transduced A2058-br melanoma brain metastasis (MBrM) cells, which had been selected for brain tropism by multiple rounds of *in vivo* passage (Fig. 1A). This methodology enables to follow the fate of each tumor cell at high temporal and spatial resolution during brain colonization^46,47^ and simultaneously allows measuring cellular Ca^2+^ fluctuations in real time (Fig. 1B) through the chronic cranial window. Using a custom developed semi-automatic imaging analysis pipeline (Fig. 1C), we quantified the dynamic Ca^2+^ activity of brain-tropic A2058-br cells (Fig. 1B). Moreover, Ca^2+^ activity was measurable in the brain-metastatic NSCLC model NCI-H1915 (Fig. 1D, Supplementary Video 1) and the patient-derived MBrM model DDMel31 (Suppl. Fig. 1A).

**Figure 1.**
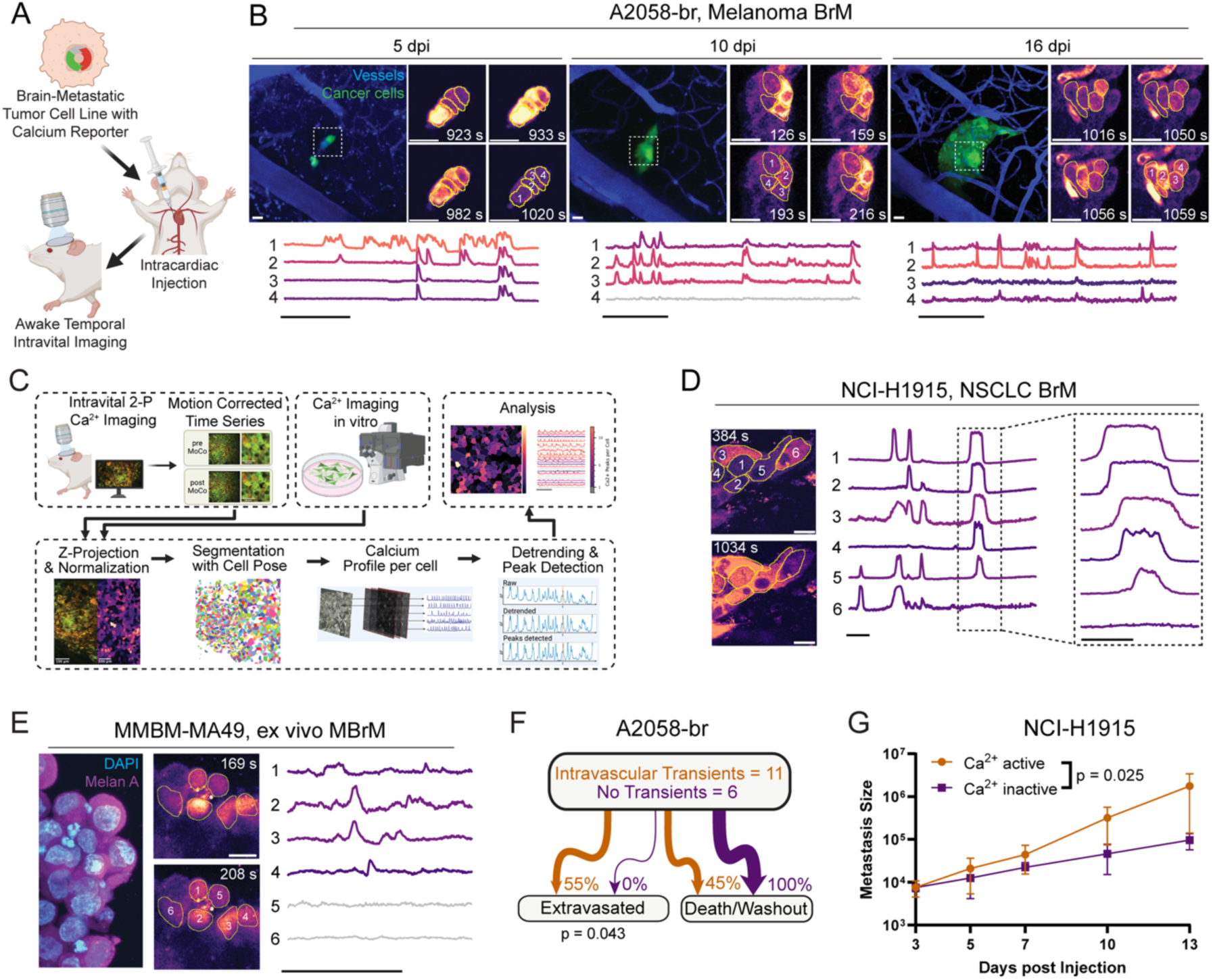
Spontaneous Ca^2+^ transients in models of brain metastasis. (A) Workflow of longitudinal in-vivo imaging of preclinical brain metastasis models in awake mice. Created in https://BioRender.com. (B) Longitudinal in vivo Ca^2+^ imaging of A2058-br brain metastases at different disease stages after intracardiac tumor injection. Top: 2D projection of the same brain region imaged at 5, 10, and 16 days post injection (dpi), showing vessels (blue) and A2058-br cancer cells (green). Dashed white square indicates the tumor region. Insets: Zoom-in on the tumor site, Sequential frames showing Ca^2+^-indicator GCaMP6s fluorescence intensity over time for the indicated regions. Scale bar = 20 µm. Bottom: Ca²⁺ traces of selected tumor cells over time. Scale bar = 5 min. (C) Workflow of custom, semi-automated image analysis and quantification for in vitro and in vivo Ca^2+^ recordings. Created in https://BioRender.com. (D) In vivo Ca^2+^imaging of NCI-H1915 brain metastasis after intracardiac injection. Left: Sequential frames of GCaMP6s fluorescence intensity over time. Scale bar = 20µm. Middle: Ca²⁺ traces of selected tumor cells over time. Scale bar = 5 min. Dashed white square indicates zoom-in. Right: Zoom-in of Ca²⁺ traces, Scale bar = 1 min. (E) In-vitro Ca^2+^ imaging of ex vivo, patient melanoma brain metastasis derived tumoroid MMBM-MA49 Left: Staining for DAPI (cyan) and Anti-Melanoma antibody (magenta). Middle: Filmstrip of chemical Ca^2+^ sensor Calbryte-520 fluorescence intensity over time. Scale bar = 20µm. Right: Ca^2+^ traces of selected cells over time. Scalebar = 5min. (F) Quantification of extravasation fate comparing intravascular cells showing no Ca^2+^ transients and those showing transients. Chi-square test, n=11/6 metastases from n=3 mice. (G) Longitudinal tumor volume of NCI-H1915 in vivo across days post heart injection. Ca^2+^ active (red; n = 6), Ca^2+^ inactive (gray; n = 4). Group means ± SEM indicated. Mixed-effects linear model tested group × day interaction on log volume, accounting for mouse and tumor variability. P-value indicates slope difference. n = 40 datapoints for n=3 mice. See also Figure S1.

To investigate if Ca^2+^ transients are (at least partially) a tumor-intrinsic property and occur independent of the brain environment, Ca^2+^ imaging of *in vitro* monocultures of the MBrM model A2058-br, the patient-derived MBrM model DDMel31, and the brain-metastatic NSCLC model NCI-H1915 was performed (Suppl. Fig. 1B-D, Supplementary Video 2). Similar to their *in vivo* counterparts, BrM cells displayed similar patterns of Ca^2+^ activity in the *in vitro* monocultures, suggesting a cancer cell-autonomous generation of these transients (Suppl. Fig. 1E-J). Additionally, *ex vivo* imaging of melanoma BrM patient-derived tumoroids cultured under stem-like conditions revealed similar patterns of complex Ca^2+^ activity (Fig. 1E), demonstrating their presence in the human disease.

Dynamic and long-term intravital microscopy demonstrated stable Ca^2+^ activity during disease progression (Suppl. Fig. 1K-L). Indeed, Ca^2+^ activity already occurred during intravascular arrest of circulating cancer cells (Suppl. Fig. 1M), demonstrating that those Ca^2+^ fluctuations occur without input from cells of the brain parenchyma, such as neurons^7,8,11,44^. Fate-mapping these cells demonstrated that only Ca^2+^-active cancer cells successfully extravasated and formed brain metastases (Fig. 1F), suggesting that intracellular Ca^2+^ transients are an intrinsic feature of brain metastasis-proficient cancer cells. Furthermore, the Ca^2+^-active, extravasated cancer cells had significantly higher growth rates compared to Ca^2+^-inactive brain metastases (Fig. 1G). Together, these findings suggest a brain seeding-promoting function of Ca^2+^ activity in BrM over multiple steps of the brain metastatic cascade.

### Collective Coordinated Ca²⁺ Activity Exists without Pacemaker Cells

In primary brain tumors, coordinated tumor cell network-wide Ca²⁺ activity patterns drive proliferation and therapy resistance^35,38^. Similarly, brain metastases displayed significant synchronicity in their Ca^2+^ activity patterns compared to a random distribution of activity across the cellular network, both *in vitro* and *in vivo* and across entities (Fig. 2A-B, Suppl. Fig. 2A-H). This co-activity between neighboring cancer cells suggests a communicating cancer cell network of the brain metastatic tumor cells across multiple extracranial entities. In glioblastoma, tumor cell networks are driven by a small subpopulation of pacemaker-like cells that activate gap junction-connected other network cells by the Ca^2+^-dependent potassium channel KCa3.1, which stabilizes Ca^2+^ oscillations, but only under distinct intracellular conditions of other Ca^2+^-relevant signaling molecules^39,48^. As KCNN4, encoding for KCa3.1, is highly expressed in preclinical BrM models (Fig. 2C), we assessed whether similar pacemaker-like cells might exist in BrM. However, no equivalent ‘pacemaker’-like periodic cell subpopulation defined by a high periodicity of multicellular Ca^2+^ oscillation was identified in BrM, in contrast to all glioblastoma models studied so far^39^ (Fig. 2D-E). Similarly, in contrast to glioblastoma, pharmacological inhibition of KCa3.1 with either TRAM34 or Senicapoc neither affected Ca^2+^ activity (Fig. 2F) nor proliferation (Fig. 2G), consistent with the absence of pacemaker-like cells in BrM. This points to relevant differences of intracellular Ca^2+^ signaling in primary and metastatic brain tumors. Taken together, BrM networks exhibit spontaneous and synchronous Ca^2+^ activity but lack the pacemaker-like cells characteristic of glioblastoma, indicating a collective multicellular generation of these oscillations that are not depending on a small subpopulation of cancer cells.

**Figure 2.**
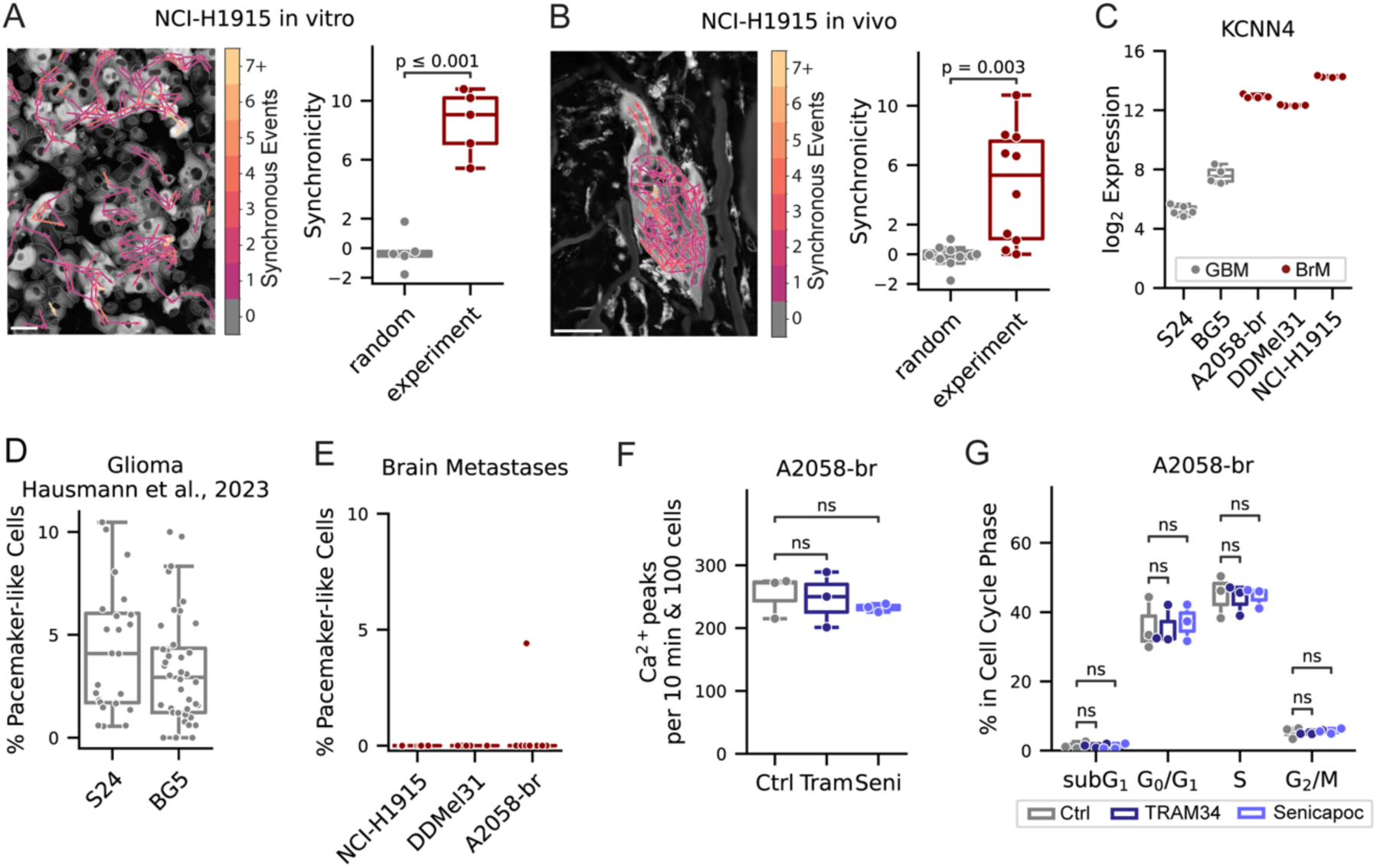
Brain metastases exhibit coordinated Ca^2+^ network activity. (A) Left: Network map of Ca^2+^ activity synchronicity in NCI-H1915 cells in vitro. Connections between neighboring cells are color-coded according to the number of synchronous Ca^2+^ events. Scale bar = 50 µm. Right: Synchronicity of Ca^2+^ activity in NCI-H1915 cells in vitro. Two-sided Welch’s t-test, n = 5 (B) Left: Network map of Ca^2+^ activity synchronicity in NCI-H1915 cells *in vivo*, 10 days post intracardiac injection. Connections between neighboring cells are color-coded according to the number of synchronous Ca^2+^ events. Scale bar = 50 µm. Right: Synchronicity of Ca^2+^ activity in NCI-H1915 cells in vivo. Two-sided Welch’s t-test, n=10 recordings from 3 mice. (C) KCNN4 Expression across glioblastoma (S24, BG5) and brain metastasis cell lines (A2058-br, DDMel31, NCI-H1915). One-way ANOVA p < 0.001; Tukey HSD p < 0.001 for all pairwise comparisons except DDMel31 vs. A2058-br (p = 0.097). (D) Percentage of pacemaker-like cells *in vivo* in Ca^2+^ recordings of patient-derived glioblastoma models S24 and BG5. Data adapted from Hausmann et al., 2023. (E) Percentage of pacemaker-like cells *in vivo* in Ca^2+^ recordings of brain metastases of lung cancer (NCI-H1915) and melanoma (DDMel31, A2058-br) models. (F) Ca^2+^ activity in A2058-br cells after treatment with 1µM TRAM34, 1µM Senicapoc or vehicle. One-way ANOVA p-value 0.723, n=3. (G) Analysis of proportion of cells in cell cycle phases (subG1, G0/G1, S, G2/M) in A2058-br cells following treatment with 1µM TRAM34, 1µM Senicapoc or vehicle. n=3. Two-way repeated-measures ANOVA with p-value for interaction between treatment and cell cycle phase p=0.9376. See also Figure S2.

### Ca^2+^ Activity Characterizes a Neuronal-like and Proliferative Cell State

To better understand the impact of Ca^2+^ oscillations on cancer cell function, we next compared brain-metastatic melanoma cells showing different levels of Ca^2+^ activity on a transcriptional level. Hereto, we utilized a recently established Ca^2+^ integrator, the CaProLa labelling system which fluorescently labels cells based on their cumulative Ca^2+^ activity over time (Fig. 3A)^49^, to separate cells into subpopulations with high, medium and low Ca^2+^ activity over time (CaProLa_5_^high^, -^medium^, and -^low^ respectively, Fig. 3B and Suppl. Fig. 3A-C). Comparing CaProLa_5_^high^, CaProLa_5_^medium^ and CaProLa_5_^low^ conditions, a distinct shift in gene expression patterns was evident (Fig. 3C-D, Suppl. Table 1). Gene set enrichment analysis revealed that the CaProLa_5_^high^ population was enriched in gene sets related to nucleotide biosynthesis, extracellular matrix (ECM) metabolism, G-protein-coupled receptor (GPCR) signaling and IGF signaling while showing a reduction in pathways associated with mitosis and vesicular transport (Fig. 3E, Suppl. Table 2). Focusing on individual genes that are differentially regulated, we identified several genes with functions implicated in neuronal cellular behavior such as synaptic plasticity (*NLGN4X*, *NTRK*, *BDNF*, *SHANK3*) and neural development (*LHX2*/*LHX6*, *PAX6*/*PAX9*) being upregulated in CaProLa_5_^high^ cancer cells, correlating Ca^2+^ activity with transcriptional activation of neuronal features (Fig. 3F). Remarkably, *FOS*, an immediate early gene (IEG) and established marker for neuronal Ca^2+^ activity^42,43^ which promotes proliferation in neural progenitor cells^50,51^, was upregulated in CaProLa_5_^high^ tumor cells (Fig. 3G). Aside from *FOS*, other IEG such as *JUN* and *ATF3*, were also upregulated in cancer cells with time-integrated high cytosolic Ca^2+^ (Fig. 3D). Gene-set enrichment analysis of a previously reported gene set^52^ validated a significant upregulation of IEGs as well as an increase of the downstream delayed primary response genes and secondary response genes, which are downstream of IEGs (Fig. 3H, Suppl. Fig. 3 D-E). The IEG machinery sits at the convergence point of multiple signaling pathways, many of whom are regulated by Ca^2+^ as a second messenger and can lead to downstream activation of cell proliferation and dedifferentiation^53,54^. We therefore analyzed the cell cycle gene expression of CaProLa_5_^high^ cells and found an overall strong correlation with gene expression during the G_1_/S phase ^55^ (Suppl. Fig. 3F). In agreement, G_1_/S pathway gene-set enrichment was observed^56^ (Fig. 3I). To validate the role of Ca^2+^ activity on cell cycle progression, we correlated *in vitro* Ca^2+^ recordings to stainings with the thymidine analogue 5-Ethynyl-2′-deoxyuridine (EdU), which is integrated into the cellular DNA during DNA replication in the S phase (Fig. 3J, Suppl. Fig. 3G). Separating cells into three distinct groups based on Ca^2+^ activity, akin to the CaProLa subpopulations, revealed that Ca^2+^ activity^low^ cells showed significantly less proliferation than the Ca^2+^ activity^medium^ and Ca^2+^ activity^high^ subpopulations (Fig. 3K, Supp. Fig. 3H). Similarly, cells labelled by EdU displayed significantly higher Ca^2+^ activity (Fig. 3L, Suppl. Fig. 3I). To summarize, Ca^2+^ transients in brain-metastatic cells correlate with the activation of cell cycle pathways and are linked to proliferation. This is consistent with the *in vivo* brain recordings where tumor growth was faster in Ca^2+^ active BrM (Fig. 1G).

**Figure 3.**
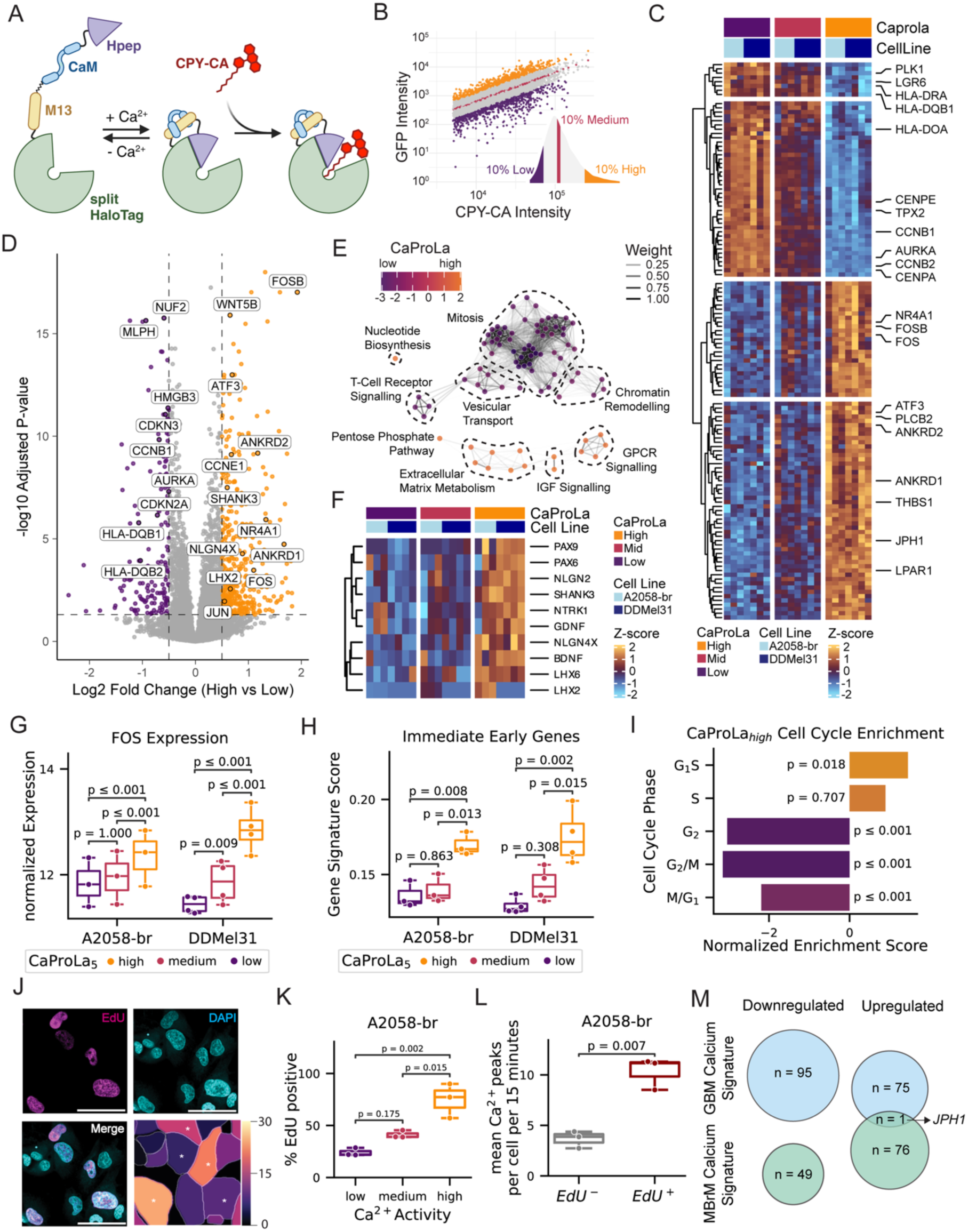
**Transcriptomic profiling of Ca^2+^ activity**. (A) Schematic of the mechanism of action of the Ca^2+^ integrator CaProLa. Created in https://BioRender.com. (B) FACS Plot of A2058-br cells expressing GFP-CaProLa5 after 30 minutes of labelling with CPY-CA dye gated into three populations, corresponding to the top 10%, medium 10% and low 10% cells based on labelling intensity. Bottom right insert: Histogram of CPY-CA / GFP ratio. (C) Heatmap showing z-score-normalized gene expression of the CaProLa signature (see methods for details) stratified by CaProLa status and cell line (A2058-br, DDMel31). Genes are hierarchically clustered. Select gene names are indicated. (D) Volcano plot showing differential expression analysis contrasting CaProLa_5_^high^ versus CaProLa_5_^medium^ and CaProLa ^low^ across the A2058-br and DDMel31 models. Genes with adjusted p-values < 0.05 and absolute log2 fold change > 0.5 are highlighted; selected genes are annotated. (E) Enrichment map of clusters of gene sets differentially regulated in CaProLa^high^ versus CaProLa^medium^ and CaProLa^low^ samples in the A2058-br and DDMel31 model (see methods for details). Common motifs in clusters were manually annotated and are highlighted in dashed circles. (F) Heatmap showing z-score transformed gene expression of neuronal genes across CaProLa status and models. (G) Boxplot of FOS expression across CaProLa status and models. DESeq2 adjusted p-values for each comparison were corrected for multiple testing using Bonferroni correction. n= 3/4 respectively. (H) Boxplot of Immediate Early Genes signature score across CaProLa status and models. Two-way ANOVA for cell line (p = 0.899) and CaProLa status (p ≤ 0.001). One-way ANOVA for CaProLa status in A2058-br (p = 0.006) and DDMel31 (p = 0.002). Post-hoc pairwise Tukey’s HSD p-values are shown in the figure. n = 3/4 respectively. (I) Gene set enrichment analysis of cell cycle signatures shows significant enrichment of G1/S and depletion of G2, G2/M, and M/G1 signatures in CaProLa^high^ cells compared to CaProLa^mid^ and CaProLa^low^. (J) Correlation of Ca^2+^ Imaging and EdU staining in A2058-br. EdU (magenta, upper left) and DAPI (cyan, upper right) staining of the same region (bottom left). Bottom right: Heatmap of Ca^2+^ peaks in 15 minutes of recording. EdU-positive nuclei are annotated with (*). Scale bar = 50µm. (K) Percentage of A2058-br EdU positive cells based on their Ca^2+^ activity status. A2058-br cells were split into Ca^2+-low^ (0-10th percentile of Ca^2+^ activity), Ca^2+-medium^ (45-55th percentile) and Ca^2+-high^ (90-100th percentile). n=3 independent recordings. Repeated-measures one-way ANOVA, 0.004. Post-hoc pairwise Tukey’s HSD p-values are shown in the figure. (L) Mean Ca^2+^ activity of A2058-br cells depending on EdU status. Two-sided Welch’s t-test, n=3 independent recordings. (M) Overlap between the MBrM Ca^2+^ Signature and the GBM Ca^2+^ Signature developed by Hai, Hoffmann et al. Nat Comms 2024. Only one gene, *JPH1*, is shared between entities and models. See also Figure S3.

We next set out to compare whether the Ca^2+^-dependent transcriptomic changes are similar or rather different between brain metastases and primary brain tumors, i.e. glioblastoma (GBM). To this end, we generated a MBrM Ca^2+^ Signature (Suppl. Fig. 3J, Suppl. Table 3) and compared it to the published GBM Ca^2+^ Signature^57^. Overlap between the signatures was non-existent except for *JPH1* (Fig. 3M), and the GBM Ca^2+^ signature did not correlate with the MBrM CaProLa^high^ state (Suppl. Fig. 3K). These findings reveal cancer cell type- and cell state-specific effects of Ca^2+^ on gene regulation and further support the fundamental differences in Ca^2+^ regulation and how it influences tumor pathobiology in primary and metastatic brain tumors.

### Molecular Mechanisms of Collective Ca^2+^ Activity

Given their association with proliferation, we set out to better understand the underlying mechanisms behind the tumor cell-intrinsic Ca^2+^ oscillations by testing the contributions of different entry routes of Ca^2+^ into the cytoplasm. For this we used an *in silico* model of Ca^2+^ regulation in non-excitable cells to demonstrate that a negative feedback loop involving inositol trisphosphate receptors (IP_3_R) in the endoplasmic reticulum (ER) and Ca^2+^ channels in the plasma membrane is sufficient to generate and sustain Ca^2+^ oscillations^58,59^ (Fig. 4A, Suppl. Fig. 4A). Fitting this model to the experimentally recorded data recapitulated the observed Ca^2+^ oscillations *in vitro* (Fig. 4B). Moreover, *in silico* inhibition of Ca^2+^ influx across the plasma membrane, modelling the effect of EGTA treatment, fully abrogated Ca^2+^ oscillations (Fig. 4C). We therefore tested experimentally whether chelation of extracellular ions with EGTA silenced Ca^2+^ oscillations in melanoma BrM cells *in vitro*, showing indeed the same effect predicted by the model (Fig. 4D, Suppl. Fig. 4B). Similarly, depletion of Ca^2+^ from ER using the SERCA-inhibitor Thapsigargin (Tha) completely abrogated Ca^2+^ activity in the model*, a*nd this was also confirmed experimentally *in vitro* (Fig. 4E, Suppl. Fig. 4C-E). This suggests that both influx of Ca^2+^ across the plasma membrane and subsequent downstream amplification via the ER are required for Ca^2+^ oscillations in cancer cells (Fig. 4F).

**Fig. 4.**
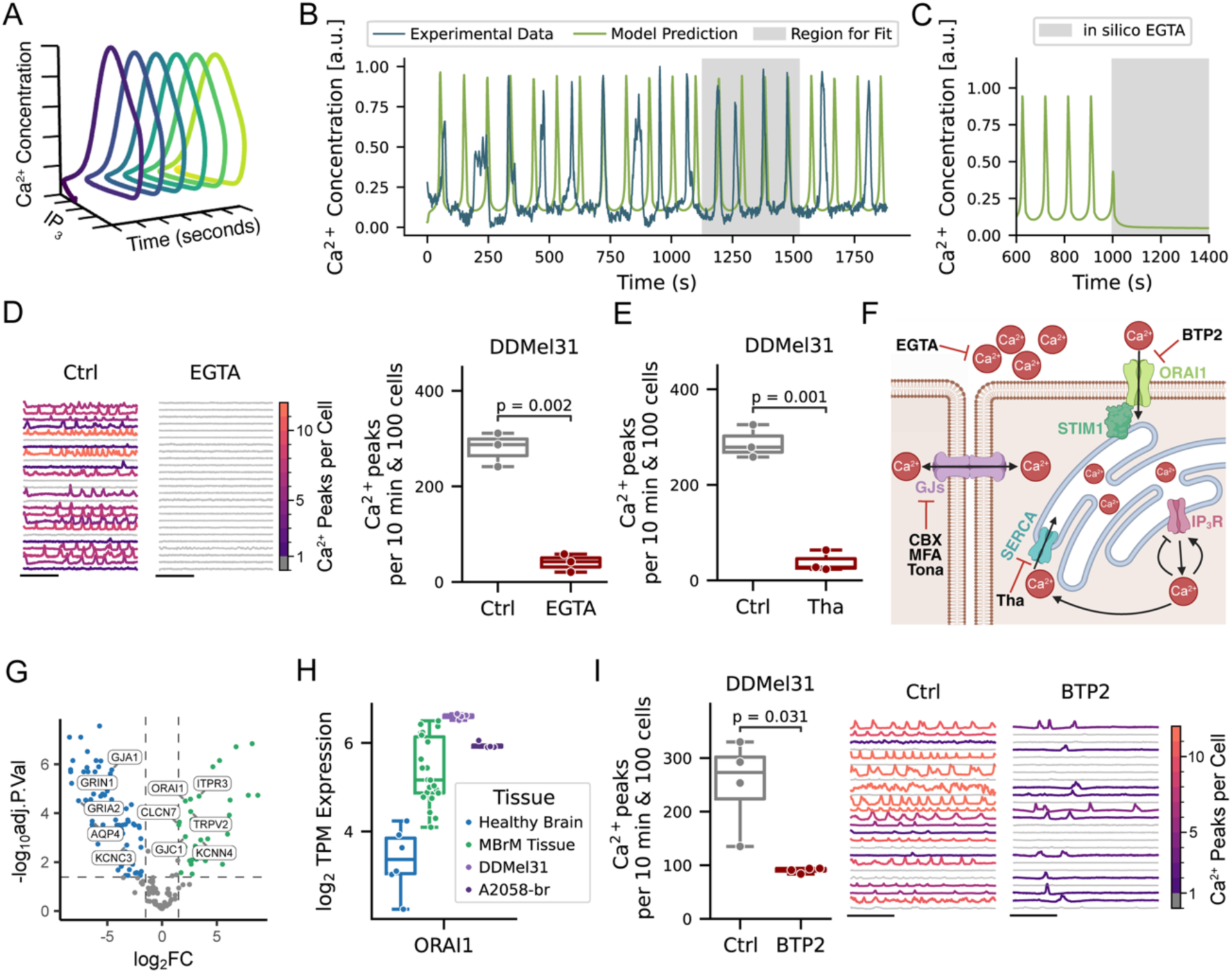
Pathways sustaining Ca^2+^ oscillations in MBrM. (A) Computational simulation of cytosolic Ca^2+^ and IP_3_ concentrations over time (see methods for details). The model demonstrates the emergence of stable Ca^2+^ oscillations. (B) Overlay of computational model (green) to experimentally recorded Ca^2+^ traces (blue) from brain-metastatic melanoma cells. Grey area indicates the time of window used for model fitting. (C) In silico simulation of Ca^2+^ dynamics after extracellular Ca^2+^ chelation with EGTA (grey area), showing rapid loss of Ca^2+^ oscillations. (D) Left: Representative Ca^2+^ traces per cell showing loss of Ca^2+^ activity after 2mM EGTA treatment. Scalebar = 5 minutes. Right: Boxplot showing Ca^2+^ activity of DDMel31 cells with vehicle and 2 mM EGTA treatment. Two-sided Welch t-test, n=3 independent recordings. (E) Boxplot showing Ca^2+^ activity of DDMel31 cells under control and 1 µM Thapsigargin (Tha) treatment. Two-sided Welch t-test, n = 3 independent recordings. (F) Schematic of the pathways involved in the generation of Ca^2+^ oscillations in MBrM. Created in https://BioRender.com (G) Volcano plot of ion channel expression in MBrM versus healthy brain tissue. x-axis: log₂ fold change; y-axis: –log₁₀ adjusted p-value. Ion channels with an adjusted-pvalue < 0.05 and absolute log2 fold change > 1 are highlighted. (H) Boxplots showing TPM-normalized gene expression (log₂ scale) of ORAI1 across healthy brain, MBrM tissue, and two MBrM cell lines (DDMel31, A2058-br). n=6 healthy brain samples, n=21 MBrM samples, n=4 DDMel31 samples, n=4 A2058-br samples. (I) Left: Boxplot showing number of Ca^2+^ peaks per 10 min per 100 DDMel31 cells treated with vehicle or 10 µM BTP2. Two-sided Welch’s t-test, n=4 independent recordings. Right: Representative Ca^2+^ traces per cell showing suppression of Ca^2+^ oscillations with BTP2 treatment. Scalebar = 5 minutes. See also Figure S4

Next, we set out to identify which ion channels are specifically upregulated in brain metastases and could mechanistically play a role in Ca^2+^ oscillations and are thus potentially amenable to therapeutic targeting. To this end, we performed an ion channel subgroup analysis of RNA sequencing data from a cohort of patient-derived BrM resections^60,61^. Compared to the transcriptome of 6 healthy brain samples within the same cohort, 30 ion channels were found to be significantly upregulated in human BrM (Fig. 4G, Suppl. Table 4). Among ion channels highly expressed in MBrM cell lines and those upregulated in human BrM cell lines is ORAI1 (Fig. 4H, Suppl. Fig. 4F). ORAI1 encodes the ion channel of the Ca^2+^ release activated Ca^2+^ (CRAC) complex^62^(Fig. 4F) and has previously been implicated in melanoma progression and invasiveness^63,64^. Indeed, in two MBrM models, the ORAI1 inhibitor BTP2 significantly reduced Ca^2+^ oscillations (Fig. 4I, Suppl. Fig. 4G). ORAI1 is expressed in most human cancers, and the mechanistical *in silico* prediction and experimentally validated necessity of ORAI1 for cytosolic Ca^2+^ oscillations in brain metastatic melanoma presented here support the clinical translation of ORAI1 targeting^65,66^.

### Gap Junction Connections Between Tumor Cells Enable Ca^2+^ Co-Activity

Gap junctions can permeate second messengers like IP3 and Ca^2+^ to neighboring cells and can thereby facilitate and sustain Ca^2+^ oscillations^67^. The gap junction protein Connexin 43 has previously been identified as the key driver of primary brain tumor network communication and subsequent progression^35,37,39^. Analysis of biopsies from a cohort of melanoma BrM patients compared to healthy brain revealed that gap junction protein Connexin 45 (*GJC1*) was specifically enriched in BrM. Gap junctions were also highly expressed in brain-metastatic melanoma cell lines (Fig. 5A). Pharmacological blockade of gap junctions with Meclofenamate (MFA), Carbenoxolone (CBX) or Tonabersat significantly reduced Ca^2+^ oscillations in both melanoma and lung cancer BrM models (Fig. 5B-C, Suppl. Fig. 5A-B, Supplementary Video 3).

**Fig. 5.**
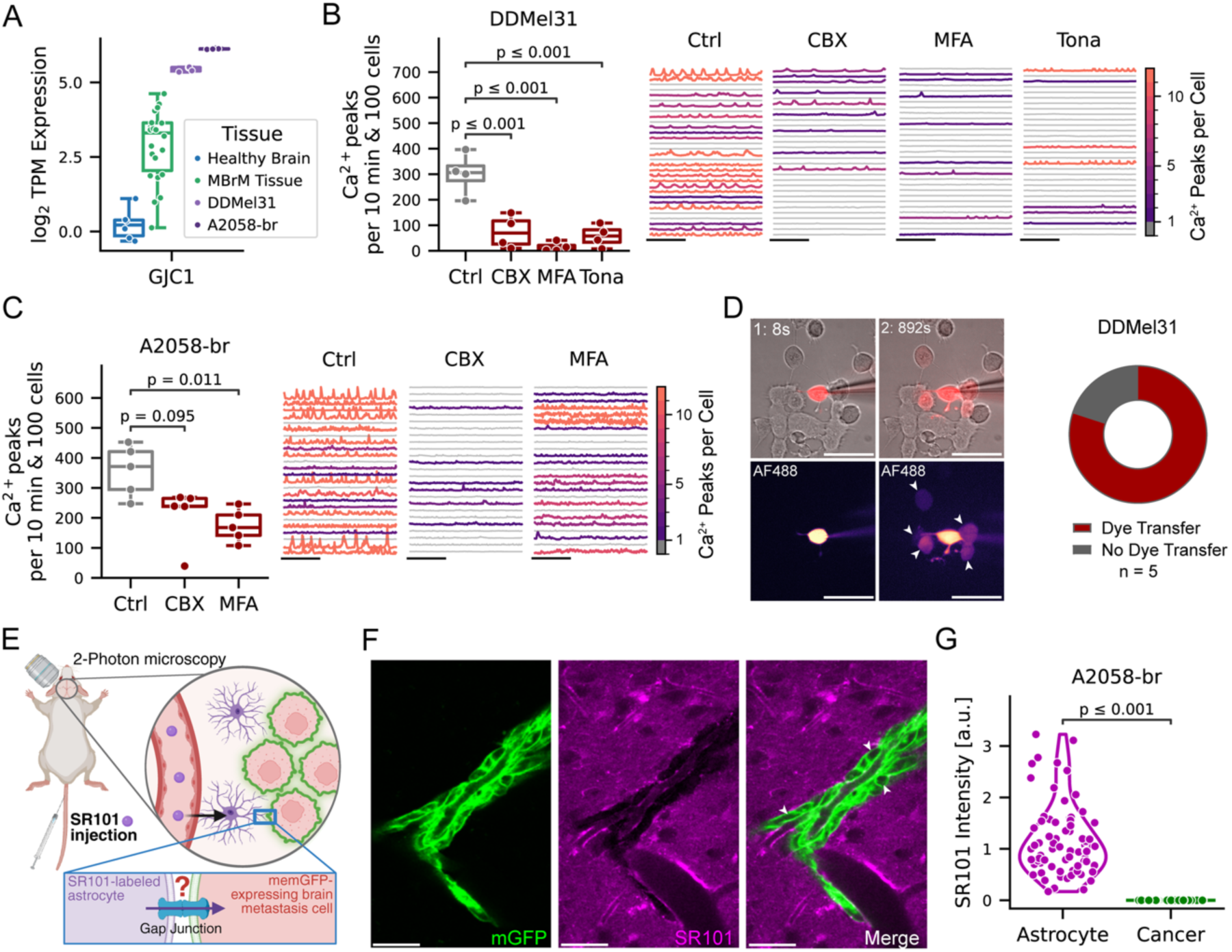
Gap-junction coupled cell signaling in brain metastases. (A) Boxplots showing TPM-normalized gene expression (log_2_ scale) of GJC1 across healthy brain, MBrM tissue, and two MBrM cell lines (DDMel31, A2058-br). n=6 healthy brain samples, n=21 MBrM samples, n=4 DDMel31 samples, n=4 A2058-br samples. One-way ANOVA p < 0.001, with post-hoc Dunn test with Holm-Sidak correction indicated in the figure. (B) Left: Ca^2+^ activity of the DDMel31 melanoma cell line after treatment with vehicle or 100 µM Carbenoxolone (CBX), 100 µM Meclofenamate (MFA) or 200 µM Tonabersat (Tona). One-way ANOVA p < 0.001 with post-hoc Dunn’s test with Holm correction indicated in figure. n=4 independent recordings. Right: Ca^2+^ traces of recordings. Cells are color-coded based on Ca^2+^ activity. Scalebar = 5 minutes. (C) Left: Ca^2+^ activity of the A2058-br melanoma cell line after treatment with vehicle or 100 µM CBX or 100 µM MFA. Kruskal-Wallis p-value = 0.012 with post-hoc Dunn test with Holm-Sidak correction indicated in the figure. n=5 independent recordings. Right: Ca^2+^ traces of recordings. Cells are color coded based on Ca^2+^ activity. Scalebar = 5 minutes. (D) Dye transfer between melanoma cells (DDMel31). Left: Representative images of a large, membrane impermeable AF488 dye injected into a single tumour cell with uptake in neighbouring cells (arrow-heads) 15 minutes later. Scalebar 50µm. Right: Quantification of observed dye transfer in n=5 experiments. (E) Schematic of SR101 astrocyte labelling mechanism. Created in https://BioRender.com (F) Representative image of astrocyte staining with SR101 (magenta) and A2058-br melanoma cell line expressing membrane-targeted GFP (green) showing no dye uptake (arrowheads). Scalebar = 50µm. (G) Quantification of SR101 fluorescence intensity of individual A2058-br metastases cells in vivo 14 days post heart injection. No uptake of SR101 in tumour cells is observed compared to astrocytes. n=72 astrocytes and n=83 cancer cells from N=2 mice. See also Figure S5.

Gap junctions enable the transfer of cytoplasmic small molecules between cells, including Ca^2+^ and other signaling molecules^68,69^. Following microinjection of a membrane-impermeable but gap junction-permeable dye into single cells, we confirmed time-dependent dye transfer to neighboring cells, indicating the presence of functional gap junction connections between tumor cells (Fig. 5D, Suppl. Fig. 5C-D). Notably, dye transfer was observed through intermediary cells, too, highlighting the multicellular interconnectivity of these tumor cell networks (Fig. 5D).

Next, we investigated whether gap junctions also engage in tumor-astrocyte gap junction coupling, as described for primary brain tumors^10,35,57,70^ and in some brain-metastatic models^31^. To investigate this, we monitored cellular SR101 dye transfer *in vivo* in real time, a robust method to detect tumor cell-astrocyte connections^10,70^. SR101 is preferentially taken up by endothelial cells and glia cells and then distributed across the astrocytic network via gap junctions^71^ (Fig. 5E, Suppl. Fig. 5E, F) and via gap junctions to tumor cells^10,35,57,70^. Although we could track uptake of the SR101 dye within the glial network and in glioblastoma cells *in vivo*, as expected (Suppl. Fig. 5G, H), such dye transfer was not observed in brain metastasis cells used in our study in the live mouse brain (Fig. 5F-G). Taken together, this makes it unlikely that functional tumor cell-astrocyte gap junction connections are operational in our experimental systems. Therapeutic inhibition of gap junctions should therefore lead to effects that are related to tumor cell-intrinsic mechanisms. In conclusion, BrM cancer cells form interconnected, homotypic networks which drive coordinated Ca^2+^ activity.

### Disruption of Ca^2+^ Activity Induces G_2_/M Cell Cycle Arrest

To investigate the relevance of BrM cancer-cell network connectivity and its Ca^2+^ activity on MBrM pathobiology, we transcriptionally profiled MBrM cells following pharmacological gap junction inhibition. In agreement with reduced Ca^2+^ activity *in vitro* (Fig. 6A), treated cells also showed significantly lower MBrM Ca^2+^ Signature scores based on transcriptomic analysis (Fig. 6B).

**Fig. 6.**
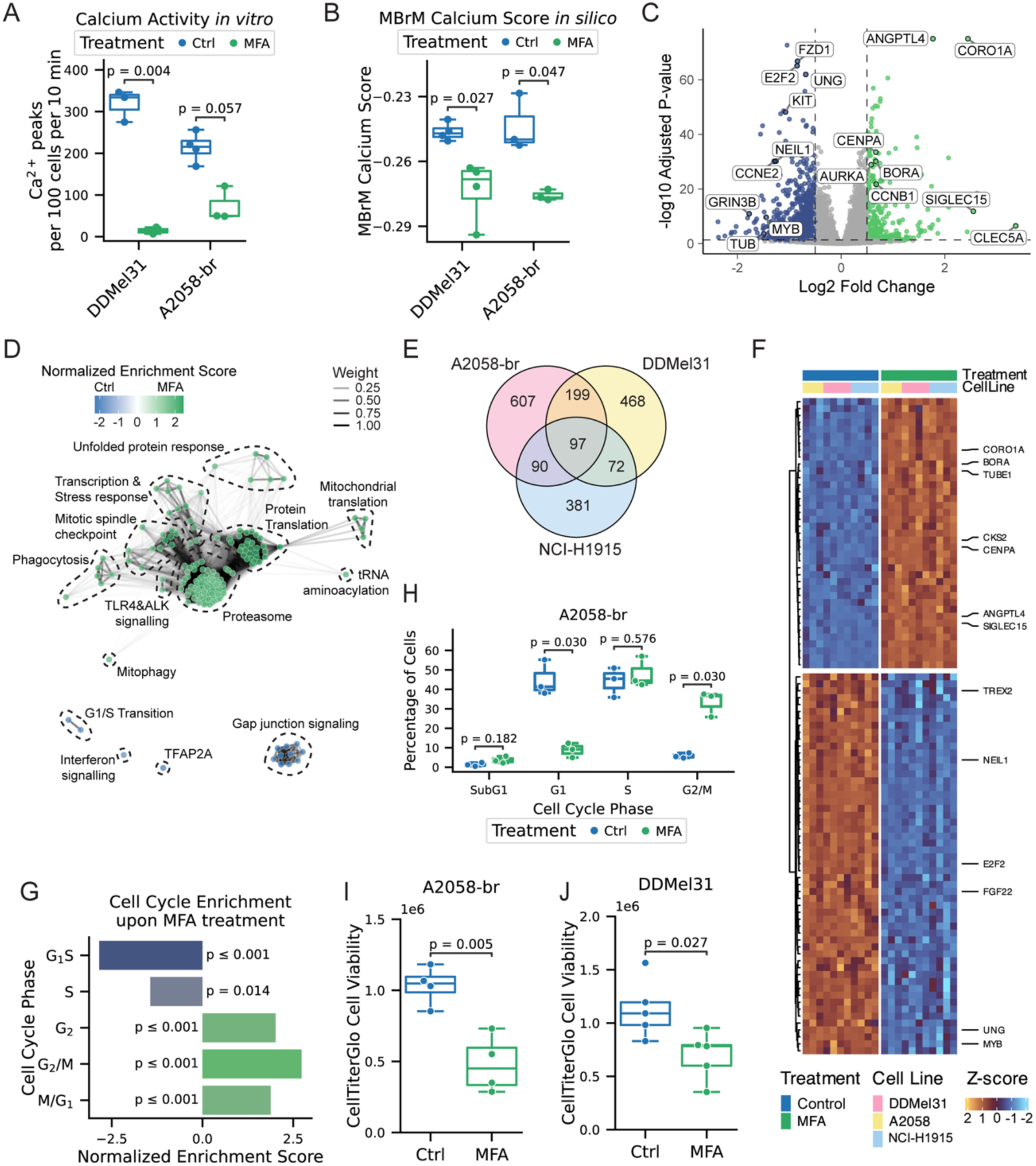
Ca2^+^ inhibition induces G_2_/M arrest and reduces cell viability. (A) Boxplots of Ca^2+^ activity following 12h treatment with Meclofenamate (MFA, 100 μM) versus vehicle control in DDMel31 and A2058-br cell lines in vitro. Two-sided Welch’s t-test, n = 3 for DDMel31. Two-sided Mann-Whitney test, n=4/3 for A2058-br. (B) Boxplot of MBrM Ca^2+^ Score in MFA-versus vehicle-treated DDMel31 and A2058-br samples. Two-sided Welch’s t-test. n = 4/3 respectively. (C) Volcano plot of RNA-sequencing differential gene expression between MFA- and vehicle-treated A2058-br and DDMel31 cells. Genes with absolute log₂ fold change > 0.5 and adjusted p-value < 0.05 are high-lighted. (D) Enrichment map of gene sets significantly up- or downregulated after MFA treatment in A2058-br and DDMel31 (see methods for details). Common motifs in clusters were manually annotated and are highlighted in dashed circles. (E) Venn diagram displaying overlap of differentially expressed genes upon MFA treatment across A2058-br, DDmel31 and NCI-H1915 in vitro. (F) Heatmap displaying z-scored variance-stabilization transformed gene expression changes of two melanoma (A2058-br, DDMel31) and one lung adenocarcinoma (NCI-H1915) cell line after 12h of treatment with 100 µM MFA. (G) Barplot showing normalized enrichment scores of cell cycle-associated gene sets from MFA-treated A2058-br, DDMel31 and NCI-H1915. FDR-adjusted p-values from gene set enrichment analysis are shown in the figure. (H) Cell Cycle Analysis of A2058-br cells after 12h of 100µM MFA treatment with quantification of subG_1_, G_1_, S and G₂/M cell fractions. Welch’s t-test or Mann-Whitney test p-values were FDR-adjusted using the Benjamini-Hochberg procedure and are reported in the figure. n= 3 independent experiments. (I) Boxplot of cell viability after 48h of vehicle versus MFA treatment in A2058-br cells, measured via CellTiterGlo assay. n = 4 independent biological experiments. Two-sided Welch’s t-test. (J) Boxplot of cell viability after 48h of vehicle versus MFA treatment in DDMel31 cells, measured via CellTiterGlo assay. n = 4 independent biological experiments. Two-sided Welch’s t-test. See also Figure S6

We identified 1131 differentially expressed genes upon gap junction inhibition (Fig. 6C, Suppl. Table 5), among them the G_2_/M markers *CENPA*, *AURKA* and *BORA* and inflammatory markers such as *ANGPTL4*, *SIGLEC15* and *CLEC5A*. Downregulated upon MFA treatment were DNA repair mechanisms (*NEIL1*, *UNG, TREX2*), the proto-oncogenes *MYB* and *KIT* as well as G1/S marker genes (*CCNE2*, *E2F2*) (Fig. 6C). In pathway analyses, as expected, a downregulation of gap junction signaling was observed (Fig. 6D). Of note, downregulation of the neural-crest lineage factor TFAP2A^72^, of G1/S transition, and of interferon gamma signaling was observed in the MFA treated group (Fig. 6D). Conversely, mitotic spindle pathways, protein degradation, and protein translation were upregulated (Fig. 6D, Suppl. Table 6). To investigate if these changes are specific to melanoma BrM or a more general feature of BrM intercellular communication, we performed transcriptional analyses of control and MFA-treated brain-metastatic NSCLC cells (NCI-H1915). Upon reduction of Ca^2+^ transients through gap-junction blockade (Suppl. Fig. 6A), the MBrM Ca^2+^ score of the NSCLC model was lowered, too (Suppl. Fig. 6B). In total, 97 genes were significantly differentially regulated across all three cell lines, NCI-H1915 and two models of melanoma brain metastasis: A2058-br and DDMel31, upon MFA treatment (Fig. 6E-F). In line with the results from the two melanoma models, cell cycle gene expression correlation analysis of NSCLC cells revealed an enrichment of G_2_/M marker genes with concordant downregulation of G1/S and S-Phase genes (Fig. 6G, Suppl. Fig. 6C). Moreover, cell cycle analysis using FACS validated the RNA sequencing results by demonstrating a striking increase in cells in the G_2_/M phase with a concordant decrease in G_1_ phase upon treatment with MFA (Fig. 6H, Suppl. Fig. 6D), indicating a G_2_/M arrest.

Taken together, gap junction inhibition induces a significant alteration in the cell cycle dynamics in a Ca^2+^-dependent manner across different BrM entities, marked by a substantial shift towards G_2_/M arrest. G_2_/M arrest has been shown to induce cell death and is the underlying mechanism of action of multiple chemotherapeutic agents ^73,74^. Indeed, after 48h of gap junction inhibition, a significant reduction in viable cells could be detected compared to control treatment in brain metastatic melanoma and lung cancer (Fig. 6I-J, Suppl. Fig. 6E). Combined, these findings demonstrate that functional disruption of gap junctions inhibits Ca^2+^ activity-driven cell cycle progression, reducing tumor cell viability.

### Ca^2+^ Co-Activity Across Cancer Entities

Having established collective Ca^2+^ signaling as targetable drivers of proliferation in melanoma and lung BrM, the question arises whether Ca^2+^ activity also plays a role in the biology of other cancer entities, or if it is a feature specific to the brain microenvironment. To this end, we examined prostate and colon cancer models, which rarely form brain metastases^75,76^. Both prostate cancer (PC3) and colon cancer (HCT116) cells exhibited coordinated Ca^2+^ transients *in vitro* (Fig. 7A, Suppl. Fig. 7A). HCT116 displayed Ca^2+^ activity frequencies comparable to brain-metastatic lung cancer and melanoma models, while PC3 showed markedly higher Ca^2+^ activity (Fig. 7B), with heterogeneous responses to gap junction inhibition (Fig. 7C, Suppl. Fig. 7B). As substantial heterogeneity in Ca²⁺ signaling activity and its mechanisms was observed both between and within cell lines (Fig. 7B, Fig. 3), we next investigated whether Ca^2+^ activity represents a transient cellular state or is rather a stable cellular feature. Following multiple rounds of CaProLa_5_-based selection (Fig. 7D), CaProLa_5_^high^ and CaProLa_5_^low^ cells partially retained their labeling intensity (Fig. 7E) and Ca^2+^ activity (Fig. 7F, Supplementary Video 4), indicating an at least partial longitudinal stability of this important cellular feature. This raises the question whether highly Ca^2+^-active cancer cells are more capable of mastering the steps of the metastatic cascade to eventually form a brain macrometastasis.

**Fig. 7.**
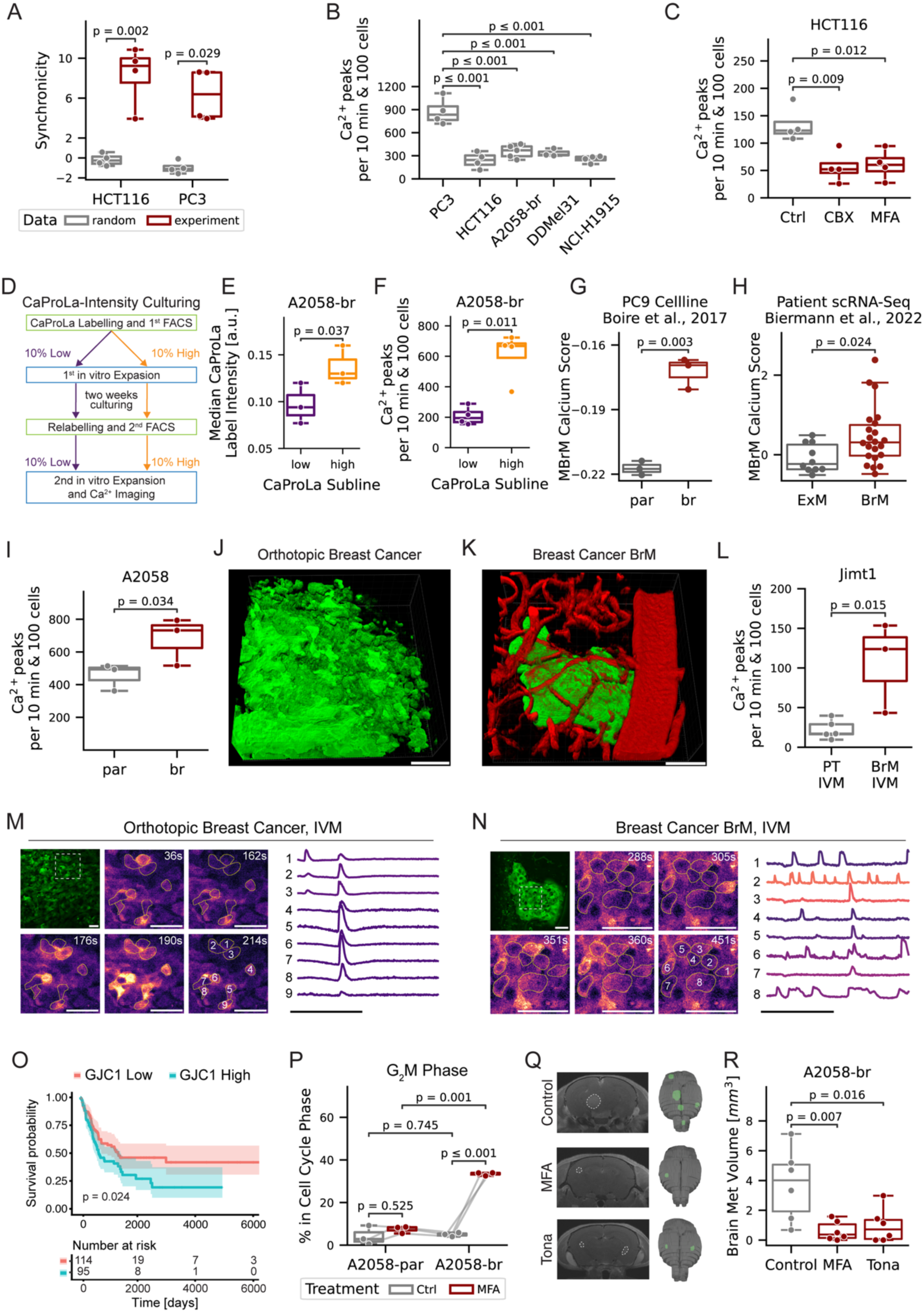
Ca2^+^ signalling is broadly conserved across cancers but intensified in metastasis. (A) Quantification of Ca^2+^ synchronicity in HCT116 and PC3 cells in vitro. Boxplots compare randomized cells with experimentally observed neighboring cells. Synchronicity is significantly higher in the experimental condition compared to random controls (n = 4, Welch’s t-Test). (B) Comparison of in-vitro Ca^2+^ activity across models. One-way ANOVA p < 0.001 with post-hoc Tukey HSD. Only significant comparisons are annotated in panel. (C) Gap junction inhibition reduces Ca^2+^ activity in HCT116. One-way ANOVA p = 0.008, Holm-corrected Dunn’s posthoc p-values indicated in the figure. n=4 independent experiments. (D) Schematic displaying the generation of CaProLa^high^ and -^low^ sublines through repeated FACS-sorting. (E) Boxplots comparing median CaProLa labelling intensity as measured by FACS between CaProLa^low^ and CaProLa^high^ sublines. n=3, two-sided Welch’s t-test. (F) Boxplots comparing Ca^2+^ activity between CaProLa^low^ and CaProLa^high^ sublines. n=4 recordings, Welch’s t-Test. (G) Comparison of in-silico MBrM Ca^2+^ signature score of the parental (par) or brain-tropic (br) PC9 lung adenocarcinoma cell line. Raw transcriptomic data obtained from GSE83132. n=3, Welch’s t-Test. (H) Comparison of in silico MBrM Ca^2+^ signature score in patient extracranial (ExM) or brain metastases (BrM). Data from GSE200218. Two-sided Mann-Whitney U test, n=10 extracranial metastases, n=21 brain metastases. (I) Comparison of Ca^2+^ activity between the A2058 parental (par) and the brain-tropic (br) subline in vitro. n=3 independent recordings, two-sided Welch’s t-test. (J) 3D reconstruction of a Jimt1 mammary fat pad tumour after orthotopic injection. Scalebar = 80µm (K) 3D reconstruction of a brain metastases after heart injection of the Jimt1 brain-tropic subline. Scalebar = 80µm (L) Comparison of Ca^2+^ activity as measured by intravital microscopy (IVM) of the orthotopic injected Jimt1 parental line (PT IVM) and of the brain metastases formed after heart injection with the brain-tropic subline (BrM IVM). n=5 recordings of N=1 mouse (PT IVM), n=3 recordings of N=3 mice (BrM IVM). Twosided Welch’s t-test. (M) In vivo Ca^2+^ imaging of breast cancer model Jimt1-par after mammary fat pad injection. Top Left: Overview of Field of View, Dashed white square indicates region of interest (ROI). Middle: Sequential frames showing GCaMP6s fluorescence intensity over time for the indicated region. Scale bar = 50 µm. Right: Representative Ca^2+^ traces of selected tumor cells over time. Scale bar = 5 min. (N) In vivo Ca^2+^ imaging of breast cancer brain metastasis model Jimt1-br after heart injection. Top Left: Overview of Field of View, Dashed white square indicates region of interest (ROI). Middle: Sequential frames showing Twitch3A fluorescence intensity over time for the indicated region. Scale bar = 50 µm. Right: Representative Ca^2+^ traces of selected tumor cells over time. Scale bar = 5 min. (O) Patient survival depending on GJC1 expression of GSE6590. n=114 (GJC1 low) versus 95 (GJC1 high). Log-rank test. (P) Quantification of A2058-par versus A2058-br cells in G_2_/M cell cycle phase of A2058-br and A2058-par under treatment with Meclofenamate (100µM) or vehicle. Two-way repeated measures ANOVA (cell line p-value = 0.008, treatment p-value = 0.005, interaction p-value = 0.005) with Holm-adjusted two-sided paired t-test p-values indicated in the figure. n=4 experiments. (Q) Representative MRI images and 3D reconstruction of brain metastases on day 28 post heart injection. Mice were treated intraperitoneally with 20 mg kg^-1^ body weight Meclofenamate (MFA), 10 mg kg^-1^ body weight Tonabersat (Tona) or vehicle. (R) Quantification of total brain metastatic burden on day 28 post heart injection measured by MRI. Oneway ANOVA p = 0.007 with post-hoc Dunnett test p-values indicated in figure. n=6 mice per group. See also Figure S7.

Consistent with this hypothesis, brain-tropic sublines derived from breast cancer (MDA-MB-231 and HCC1954) and lung cancer (PC9) exhibited significantly higher Ca^2+^ transcriptomic signatures compared to their non-brain-metastatic parental counterparts^77^ (Fig. 7G, Suppl. Fig. 7C-D). Moreover, patient-derived melanoma brain metastases displayed elevated Ca^2+^-transcriptomic signatures relative to extracranial metastatic lesions^26^ (Fig. 7H). Functional imaging corroborated these findings, as the brain-metastatic subclone of the A2058 melanoma model exhibited increased Ca^2+^ activity compared to its parental line (Fig. 7I). Furthermore, intravital imaging revealed higher Ca^2+^ activity in brain metastases formed by the Jimt1 breast cancer BrM subline compared to the Jimt-1 parental line implanted in the mammary fat pad (Fig. 7J-N, Supplementary Video 5). In line, brain-tropic cancer cells demonstrated increased expression of the gap junction protein GJC1 compared to non-brain-tropic counterparts (Suppl. Fig. 7E-G). Notably, elevated GJC1 expression was also associated with reduced overall survival in melanoma patients ^78^ (Fig. 7O). These results indicate that elevated Ca^2+^ activity and Ca^2+^ co-activity-related transcriptomic signatures are a feature of brain metastases.

### Collective Ca^2+^ Activity is a Therapeutically Targetable Feature of BrM

Blocking Ca^2+^ co-activity with Meclofenamate (MFA, Suppl. Fig. 7H) induced G_2_/M arrest in the brain-metastatic subline more than in its parental counterpart (Fig. 7P), providing further evidence for a metastasis-specific susceptibility towards targeting of Ca^2+^ communication. Therefore, inhibition of Ca^2+^ communication *in vivo* by injecting mice intracardially with A2058-br cells and treating them with the gap junction inhibitors MFA, tonabersat, or vehicle control was tested. Cranial MRI revealed a significant reduction in total intracranial tumor burden and the number of BrM in both treatment groups with the gap junction inhibitors, compared to the control group (Fig. 7Q-R, Suppl. Fig. 7I-J).

## Discussion

Here, we show that collective, coordinated Ca^2+^ activity in cancer-cell intrinsic, gap junction-coupled networks is a driver of brain metastasis and linked to neural cellular programs. These findings extend the concept of cancer-intrinsic networks and collective Ca^2+^ activity as a cancer cell-intrinsic neural feature beyond primary brain tumors^5,9,10,12,33–35,38,39,70^. In contrast to primary brain tumors, BrM generate Ca^2+^ co-activity and oscillations without cellular pacemakers, but rather in a collective manner.

In the developing brain, spontaneous intercellular Ca^2+^ waves, in part mediated by gap junctions^79,80^, drive cell cycle progression and proliferation of radial glia cells and neural progenitors^81–83^. Similarly, glioblastoma tumor cell networks display autonomous rhythmic Ca^2+^ activity that drives tumor growth^10,35,39^, suggesting that both primary brain tumors and, as we now demonstrate, brain metastases of non-neural origin hijack neurodevelopmental Ca^2+^ signaling principles to fuel malignant progression.

These Ca^2+^ oscillations are distinct from a general increase in cytoplasmic Ca^2+^. Indeed, sustained high concentrations of Ca^2+^ are often pro-apoptotic^84^, whereas oscillatory Ca^2+^ signals can regulate signaling pathways by activating NFAT^85,86^, NF-kB^85^, MAPK^87^, as well as CAMKII^88,89^ in a frequency-dependent manner ^90^.

Crucially, several of these frequency-decoded effectors have established roles not only in primary tumor biology but specifically in metastatic progression^91,92^, as they mediate migration, invasion, and metastasis formation^63,93–95^. Furthermore, metastatic cancer cell lines have been reported to display higher levels of Ca^2+^ oscillations compared to non-metastatic cell lines^96^.

The work presented here demonstrates that cancer-cell intrinsic Ca^2+^ co-activity is detectable in cells from extracranial cancers and is enriched in BrM. Brain-metastatic sublines across multiple cancer entities exhibited elevated Ca^2+^-transcriptional signatures. Additionally, brain metastases cell lines had increased expression of gap junction components, which correlate with poor patient outcome. These results suggest that heightened Ca^2+^ network activity is a specific trait that is selected for during metastasis, and at the same time a broadly conserved feature of malignancy, making it a potential therapeutic target against metastatic tumor growth.

Pharmaceutical disruption of gap junctions inhibited Ca^2+^ network communication, induced G_2_/M cell cycle arrest in both melanoma and lung cancer BrM and led to a significant reduction in intracranial metastatic burden in vivo. This demonstrates that Ca^2+^ oscillations are not only a driver of tumor proliferation but also pose an actionable vulnerability in BrM. Indeed, the gap junction inhibitor MFA is currently being evaluated in a Phase II clinical trial for glioblastoma (MecMeth/NOA-24)^97^, underlining the clinical feasibility of targeting gap-junction coupled networks in brain tumor patients. While gap junction-coupled Ca^2+^ waves are essential for tissue architecture and neurogenesis during development^98–100^, gap junction coupling in the healthy adult brain is more restricted^101,102^. In addition, heterotypical connexin 43-connected networks of BrM carcinoma cells with astrocytes can support growth in other BrM models^31^. While not observed with the cell lines used in our study, those cancer cell-astrocyte gap junction connections can in principle provide an additive therapeutic effect of gap junction inhibitors. This suggests a therapeutic window in which disrupting tumor cell network communication by gap junction inhibition disproportionately affects malignant cells - while sparing normal tissue.

## Limitations of the study

In this work, we demonstrated conserved principles of collective Ca^2+^ signaling across multiple preclinical models and patient-derived samples, while mechanistic analyses were primarily performed in melanoma and lung cancer BrM. If and how these findings can be generalized to the metastatic potential of other tumor entities, and potentially other metastatic sites, needs to be determined. Moreover, although the effect of gap junction inhibition showed clear antiCa^2+^ and anti-metastatic effects, the dynamic interplay between metastatic Ca^2+^ signaling, neural activity, and immune components of the brain microenvironment was not systematically addressed. Future studies should aim to delineate if perisynaptic^44^ as well as direct synaptic neuron-cancer crosstalk^7,8,11^ on one hand and tumor-intrinsic Ca^2+^ signaling on the other converge on the same (or similar) cellular pathways, and whether targeting both tumor-intrinsic and neuronal-induced Ca^2+^ activity might offer a synergistic approach for new neuroscience-instructed cancer therapies.

In conclusion, these findings suggest cancer-intrinsic neural features as prime drivers of metastatic colonization. Brain metastases deploy cancer-cell intrinsic Ca^2+^ network communication, which is both strongly coupled to proliferative cell-cycle programs and the adoption of a neuronal-like transcriptomic phenotype. Targeting this signaling axis exposes a vulnerability of metastatic malignant disease, offering new conceptual and therapeutic approaches in treating metastases to the brain – and potentially to other organs.

## Resource availability

### Lead contact

Further information and requests for resources and reagents should be directed to and will be fulfilled by the Lead Contact, Matthia A. Karreman.

### Materials availability

Primary cell lines, plasmids, and transduced cell lines are available upon reasonable request. Requests for resources and reagents should be directed to the lead contact.

### Data and code availability

All sequencing data will be deposited to EGA prior to publication. Analysis code and raw counts are publicly available at https://github.com/Winklerlab-HD/CaComm-BrM. The calcium imaging analysis software pyKrait is available as a python package via https://pypi.org/project/pykrait/ and the source code is available at https://github.com/Winklerlab-HD/pykrait.

## Acknowledgements

We thank the German Cancer Research Center (DKFZ) Core Facility services provided by D. Krunic, M. Brom and F. Bestvater of the Light Microscopy Core Facility, F. Blum, K. Hexel and S. Schmitt of the Flow Cytometry Core Facility, the Next Generation Sequencing Facility, the Omics IT and Data Management Core Facility (ODCF) as well as K. Dell, M. Kempfert, A. Riedasch and the Animal Caretaker Staff of the Center for Preclinical Research (ZPF). We thank Dr. Tim Holland-Letz for biostatistical support. We thank Manuel Fischer for performing MRI measurements.

This work was supported by the Bundesministerium für Forschung, Technologie und Raumfahrt (BMFTR) within the framework of the e:Med research and funding concept (01ZX1913B to M.S., 01ZX1913A to D.W. and 01ZX1913D to M.A.K.). Support from the Deutsche Forschungsgemeinschaft (DFG, German Research Foundation), project ID 259332240/RTG 2099 was addressed to F.W. and M.A.K., and project ID 404521405, SFB 1389, UNITE Glioblastoma, was addressed to D.C.H., M.R., M.O.B., W.W., F.W. and M.A.K. D.C.H. received a scholarship from the Medical Scientist Program of the Medical Faculty at Heidelberg University. M.O.B. was supported by the Emmy Noether program of the DFG (DFG, BR 6153/1-1), the Chica and Heinz Schaller Foundation and the Else Kröner-Fresenius Stiftung (2017-A25;575 2019_EKMS.23; Else Kröner Clinician Scientist professorship, 2024_EKCS.06). Further support came from the European Center for Neurooncology (EZN)’s funding line of the Dietmar Hopp foundation to F.W. and M.A.K., and by a grant from the Hertie Foundation for Excellence in Clinical Neuroscience to M.A.K. and D.H.

## Author contributions

N.R.H., N.O., F.W. and M.A.K. conceptualized this project. Investigation was performed by N.R.H., N.O., J.S., C.D.M., J.H., T.K., S.B., C.T., C.N., N.E.H., D.C.H., A.K., A.F.V., C.Z.L., Y.Y., D.D.A., V.V., D.H., M.O.B. and M.A.K. Formal analysis was performed by N.R.H., N.O. and J.S. under the supervision, administration and funding of W.W., F.W. and M.A.K. Custom code was designed and contributed to by N.R.H., N.O., S.B., A.F.V. and D.H. New methods and models were developed by N.R.H., N.O., J.S., S.B., C.N., N.P., C.Z.L., M.R. and A.F.V. Resources were provided by M.R., N.P., K.J., M.S., F.M. D.W., M.O.B., J.T., W.W., F.W. and M.A.K. The original draft was written by N.R.H., N.O. and M.A.K. with visualizations provided by N.R.H., N.O. and J.S. Review and revision by F.W. and M.A.K. All authors have substantively revised the paper and approved the submitted version.

## Methods

### Key resources table

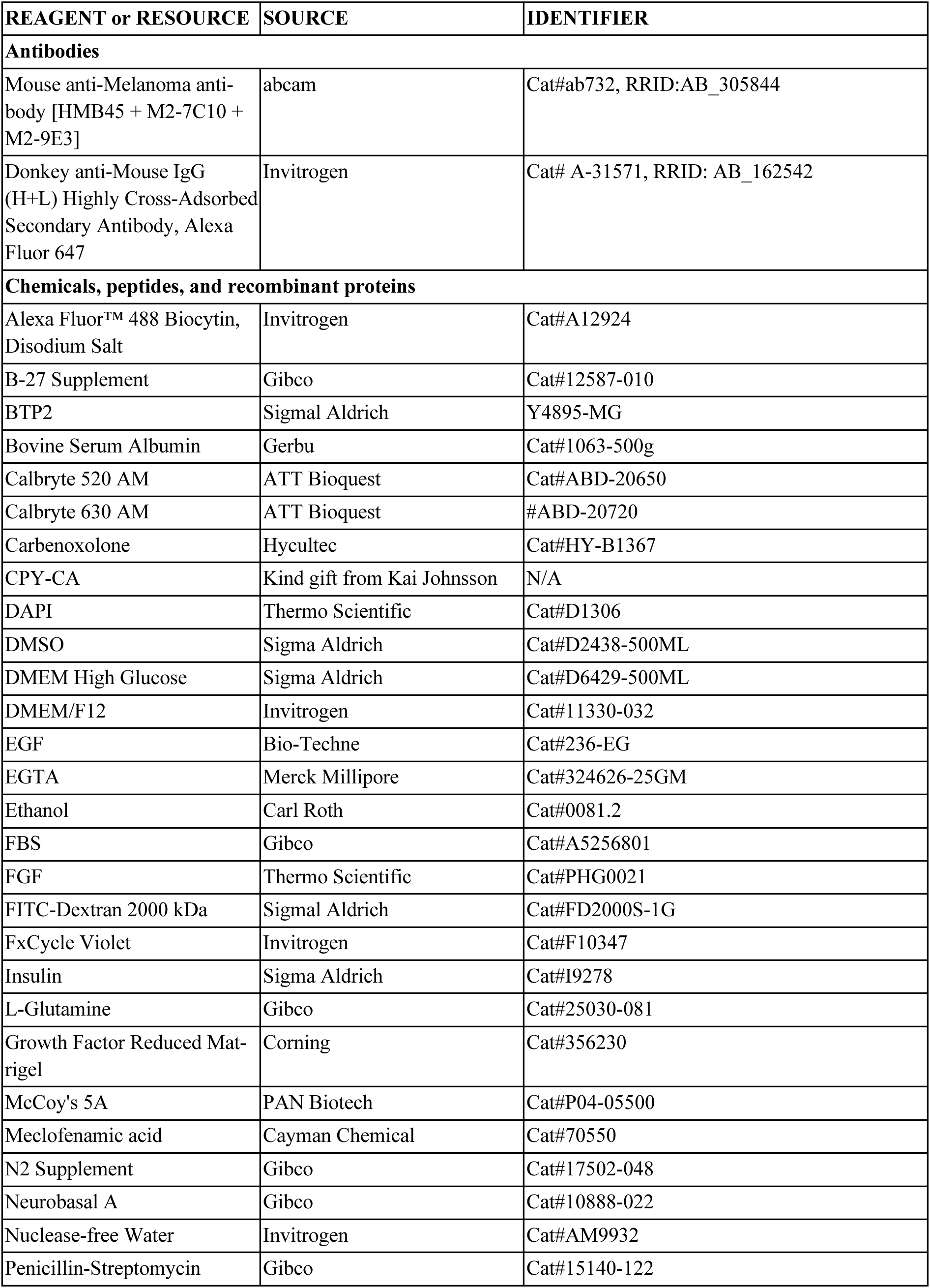
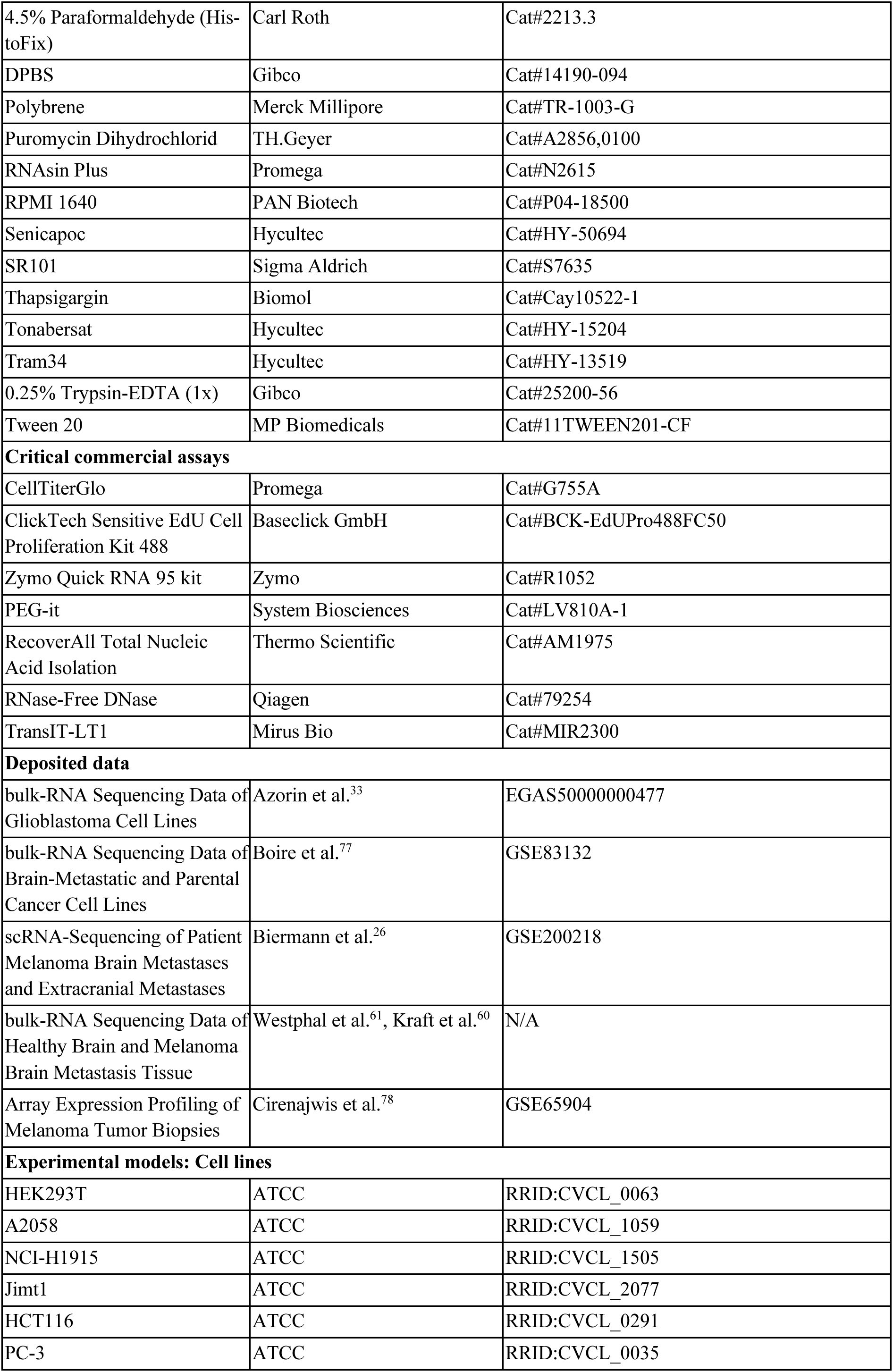
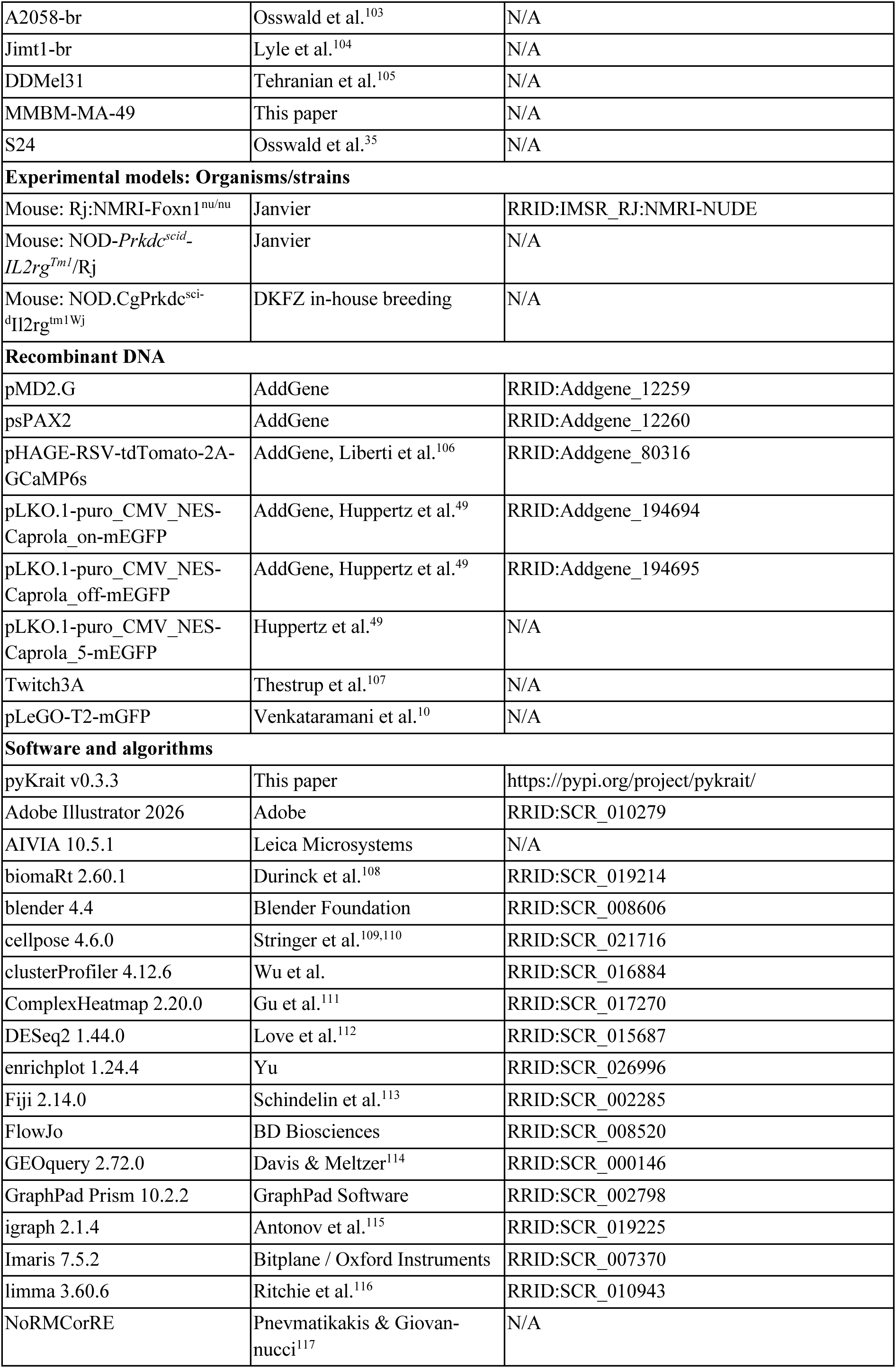
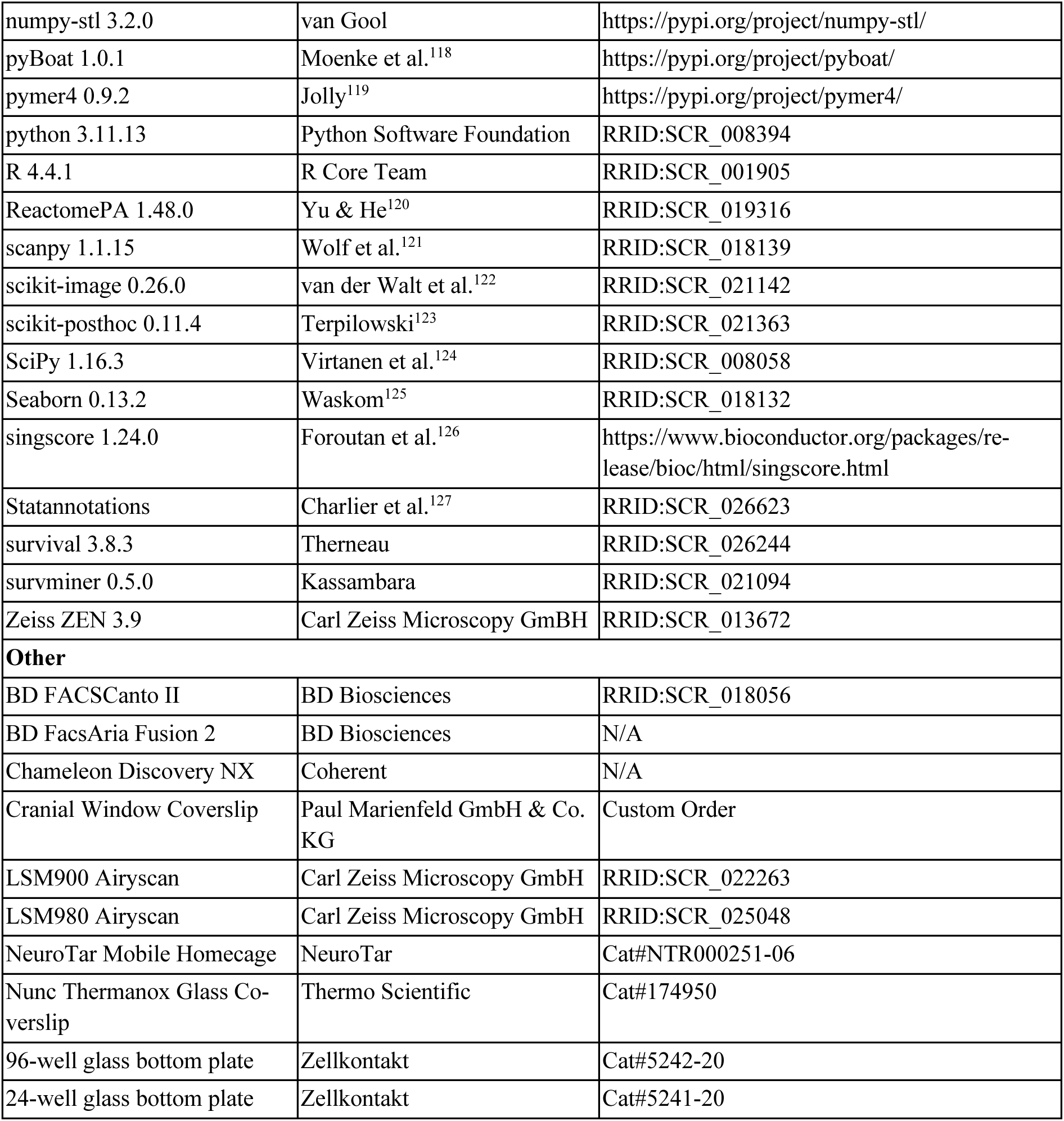

### Experimental model and study participant details

#### Mice

Immunodeficient Rj:NMRI-Foxn1^nu/nu^ (NMRI nude, Janvier Labs, RRID:IMSR_RJ:NMRI-NUDE), NOD.CgPrkdc^scid^Il2rg^tm1Wj^ (NSG, DKFZ in-house) or NOD-Prkdc^scid^-IL2rg^Tm1^/Rj (NXG, Janvier Labs) mice, aged at least 8 weeks, were used. Animals were housed in pathogen-free conditions under a 12 h light/dark cycle with ad libitum access to food and water. Male mice were used for melanoma and lung cancer studies, female mice were used for breast cancer studies. Littermates of the same sex were randomly assigned to experimental groups. All animal procedures were performed in accordance with the institutional laboratory animal research guidelines after approval of the local governmental Animal Care and Use Committee (Regional Council Karlsruhe, Germany, 359185.81/G-220/16, 35-9185.81/G-163/22, 35-9185.81/G-110/21, 35-9185.81/G-111/21, 35-9185.81/G-60/22, 35-9185.81/G-273/19). Efforts were made to minimize animal suffering and to reduce the number of animals used according to the 3Rs principles. All mice were routinely checked for clinical endpoint criteria.

#### Cancer Cell Culture

A2058 (RRID:CVCL_1059), Jimt-1 (RRID:CVCL_2077), HCT116 (RRID:CVCL_0291), PC-3 (RRID:CVCL_0035) and NCI-H1915 (RRID:CVCL_1505) were obtained from ATCC. The Jimt1-br originated from RRID:CVCL_2077 through repeat brain passaging and was a kind gift from Patricia Steeg^104^. The A2058-br cell line originated from RRID:CVCL_1059, was developed through repeat brain passaging and described previously^103,105^. The DDMel31 cell line (BRAFV600E-/-, NRAS Q61+) was isolated from a patient with a melanoma brain metastasis under ethics approval BO-EK-161032021. The MMBM-MA49 cell line was isolated from a patient with a melanoma brain metastasis under ethics approval 2018-614N-MA, 2024-640. The S24 glioblastoma spheroid model was described previously^35^.

All tumor cell lines were cultured under standard conditions (37°C, 5% CO_2_, humidified incubator). A2058, A2058-br, Jimt1 and Jimt1- br were cultured in DMEM-6429 (Sigma Aldrich, Cat#D6429-500ML) supplemented with 10% FBS (Gibco, cat. #A5256801) and 1% penicillin-streptomycin (Gibco, Cat#15140-122). DDMel31 was cultured in RPMI-1640 (PAN Biotech, Cat#P04-18500), supplemented with 10% FBS (Gibco, A5256801), 1% L-Glutamine (Gibco, Cat#25030081), and 1% penicillin-streptomycin (Gibco, 15140-122). PC3 and NCI-H1915 were cultured in RPMI-1640 (PAN Biotech, Cat#P04-18500), supplemented with 10% FBS (gibco, Cat#A5256801) and 1% penicillin-streptomycin (Gibco, Cat#15140-122). HCT116 was cultured in McCoy’s 5A media (PAN Biotech, Cat#P04-05500) supplemented with 10% FBS (Gibco, Cat#A5256801) and 1% penicillin-streptomycin (Gibco, Cat#15140-122). MMBM-MA49 was cultured in Neurobasal A (Gibco, Cat#10888-022) supplemented with 1% B27 (Gibco, cat. #12587-010), 0.5% N2 (Gibco, Cat#17502-048), 1% L-Glutamine (Gibco, Cat#25030081) while EGF (Bio-Techne, Cat#236-EG) and FGF (Thermo Scientific, Cat#PHG0021) were added to the culture flasks freshly for a final concentration of 20 ng/ml. S24 cells were cultivated in DMEM/F12 (Invitrogen, Cat#11330-032) supplemented with B27 (Gibco, Cat#12587-010), 5µg/ml insulin (Sigma-Aldrich, Cat #I9278), with EGF (Bio-Techne, Cat#236-EG) and FGF (Thermo Scientific, Cat#PHG0021) added to the culture flasks freshly for a final concentration of 20 ng/ml.

All non-primary human cell lines were authenticated based on Single Nucleotide Polymorphism (SNP) typing. During the whole duration of the study, all cell lines were tested via PCR every three months for mycoplasma contaminations and were grown on tissue-culture treated plasticware (Greiner, Cat#658175) or as indicated in the respective experiments.

### Method details

#### Virus production

Lentiviral particles were produced by co-transfecting pMD2.G and psPAX2 with the corresponding construct in HEK293T cells using TransIT-LT1 (Mirus Bio, MIR2300). pMD2.G was a gift from Didier Trono (Addgene plasmid #12259; http://n2t.net/addgene:12259; RRID:Addgene_12259). psPAX2 was a gift from Didier Trono (Addgene plasmid #12260; http://n2t.net/addgene:12260; RRID:Addgene_12260). pHAGE-RSV-tdTomato-2A-GCaMP6s was a gift from Darrell Kotton^106^ (Addgene plasmid #80316; http://n2t.net/addgene:80316; RRID:Addgene_80316). The Twitch3A calcium biosensor was a kind gift from Olga Garatschuk and Oliver Griesbeck^107^. The CaProLa calcium integrator plasmids pLKO.1-puro_CMV_NES-Caprola_on-mEGFP (Addgene plasmid #194694; http://n2t.net/addgene:194694; RRID:Addgene_194694), pLKO.1-puro_CMV_NES-Caprola_off-mEGFP (Addgene plasmid #194695; http://n2t.net/addgene:194695; RRID:Addgene_194695) and pLKO.1-puro_CMV_NES-Caprola_5-mEGFP were gifts from Kai Johnsson^49^. pLeGO-T2-mGFP was a gift from Varun Venkataramani, previously described^10^ and developed based on the mGFP sequence by De Paola et al^128^. 48h and 96h after transfection, virus was harvested using PEG-it (System Biosciences, cat. #LV810A-1) following the manufacturer’s instructions.

#### Lentiviral Transductions of Cancer Cell Lines

300 000 tumor cells were seeded in a 6-well plate (Faust Lab Science, Cat#TPP92006). 24 hours after seeding, cells were incubated with 8 ng/ml Polybrene (Merck Millipore, Cat#TR-1003-G) and 10 µl of lentivirus. Transduced cancer cells were selected either based on endogenous fluorescence by FACS sorting on a BD FACSAria Fusion, via antibiotic resistance with 1µg/ml puromycin (Th.Geyer, Cat#A2856,0100, or a combination of both.

#### Cranial Window Surgery

Cranial window surgery was performed as previously described^129^. Briefly, mice were anesthetized with Ketamine/Xylazine and transferred to a heating mat. A skin incision was made, skull bone and dura mater were removed, and a cranial window (Paul Marienfeld GmbH & Co. KG, Cat#0110990091) together with a custom titanium ring was implanted, allowing for pain-free fixation for imaging after surgery.

#### Mammary Window Surgery

Mammary imaging window (MIW) implantation was performed at least one week after orthotopic tumour implantation following a published protocol^130^ with minor adaptations. Flexible PDMS-based MIWs were obtained commercially (infenx, France). Briefly, mice were anesthetized with Ketamine/Xylazine and a ∼10 mm skin incision was made to create an implantation pocket exposing the growing tumor. The PDMS ring was inserted, unfolded beneath the skin, and skin edges placed into the groove, eliminating the need for sutures. Trapped air or accumulating debris under the window was removed by flushing with sterile saline before imaging sessions. Postoperative analgesia and prophylactic antibiotics were administered, and mice were monitored until full recovery.

#### Cancer Cell Injections

Intracardiac cancer cell injection was performed at the earliest 14 days after cranial window surgery. Mice were anesthetized with Ketamine/Xylazine. Cell lines A2058-br, NCI-H1915 and Jimt1-br (500 000 tumor cells in 100µl sterile PBS) were injected into the left ventricle of the heart. Orthotopic implantation of DDMel31 and S24 cells into the brain was performed in mice anaesthetized with Ketamine/Xylazine. The cranial window was carefully removed, the brain exposed and 100 000 cells suspended in 1µl of sterile PBS were stereotactically injected into the brain cortex at a depth of 200µm. The brain was re-sealed with a fresh cranial window (Paul Marienfeld GmbH & Co. KG, Cat #0110990091). Orthotopic implantation of 300 000 Jimt1 cells into the fourth inguinal mammary fat pad was performed using a 1:1 mixture of PBS and growth factor–reduced matrigel (Corning, Cat #356230).

#### Intravital Two-Photon Microscopy

To visualize the cerebral angiogram, mice were injected with FITC-Dextran (10mg/ml, Sigma Aldrich, Cat#FD2000S-1G) in the lateral tail vein. For awake imaging, mice were transferred to a NeuroTar mobile homecage (NeuroTar, Cat#NTR000251-06) which allows for free movement on an air-lifted treadmill. For imaging under isoflurane narcosis, mice were anesthetized with 0.5-2.0% isoflurane and transferred to a heating mat with a custom narcosis mask. Mice were placed under a ZEISS LSM980 microscope equipped with a Chameleon Discovery NX laser (Coherent). Images were acquired using a W Plan-Apochromat 20x/1.0 DIC D=0.17 M27 75mm objective (ZEISS). The cerebral angiogram was used to reliably follow up individual brain metastasis cells for weeks. TWITCH3A was visualized by exciting CFP at 860 nm, with CFP and mVenus fluorescence detected through band pass filters at 460–500 nm and 525–560 nm. GCaMP6s and tdTomato fluorophores were excited with 960nm and recorded with a nosepiece 2 channel GaAsP NDD detector (ZEISS) with Band-pass emission filters of 505–545 nm for GCaMP6s and FITC and 575-605 nm for tdTomato. Calcium imaging was performed with a sampling rate between 0.9s and 2.0s. Laser power and gain were adapted based on imaging depth to avoid phototoxicity.

#### Analysis of Brain Metastasis Volume

For quantification of the presence of Ca^2+^ oscillations in metastases across the metastatic cascade, metastases were considered to show calcium oscillations if one or more Ca^2+^ imaging time lapses demonstrated Ca^2+^ transients. To quantify the volume of the in vivo metastases, z-stack files were loaded into Imaris (Version 7.5.2, RRID:SCR_007370). The “volume” tool was used to measure the total volume of the tdTomato/red channel representing a pan cellular staining. Thresholding was kept at the suggested value for each image. Distant cells and artifacts not belonging to the main tumor mass were excluded. Volumes of cell clusters making up the metastasis were added up. For metastasis with a weak or bleached red channel, the original z-stack was loaded into Fiji ImageJ2 (Version 2.14.0/1.54f, RRID:SCR_002285) and the channels were split into red (tdTomato) and green (GCaMP6s) using the “Split channels” tool. The resulting green channel image was then subtracted from the red channel image to obtain the cleaned pan cellular red channel. The resulting image was then loaded into Imaris (Version 7.5.2, RRID:SCR_007370), original image dimensions were restored and matched to the raw file. The volume analysis was then done as described above. To compare growth trajectories between Ca^2+^-active and Ca^2+^-inactive metastases, volumes were log-transformed and a linear mixed-effects model was fitted using the lme4 framework via pymer4^119^, with group, time (day), and their interaction as fixed effects, with random intercepts for mouse and tumor identity. The interaction term (group × day) was used to test for a difference in growth slopes between groups

#### Extravasation Analysis

Individual tumor cells were tracked using intravital microscopy over multiple weeks following intracardiac injection. For quantification of the relationship between Ca^2+^ oscillations and successful extravasation, only tumor cells already arrested in microcapillaries on day 3 post injection were considered to avoid bias. The cerebral angiogram was used to reliably identify the same tumor cells and assess their respective fates (successful extravasation or death / washout).

#### SR101 Dye Transfer and Quantification

Window-bearing animals were injected with 500 000 A2058-br cells, expressing GFP in the cellular plasma membrane (mGFP), into the lateral ventricle of the heart, as described above.

2 hours before intravital two-photon microscopy, 20mg/kg SR101 (Sigma Aldrich, Cat#SHBP7695) dissolved in PBS was injected intravenously into the tail vein of tumor-bearing mice. SR101 and mGFP were excited with 960nm and emission recorded using Band-pass emission filters of 505–545 nm for mGFP and 575-605 nm for SR101. For quantification of the SR101 signal intensity, raw microscopy data was linearly unmixed using the Unmix function of ZEN 3.9 (RRID:SCR_013672) with the Automatic Component Extraction (ACE) algorithm to identify spectral overlap. Background was removed by subtracting a filtered image of the raw data (Gaussian blur with a sigma of 40 µm). Tumor Cells were segmented manually in Cellpose based on endogenous mGFP expression. SR101 positive and mGFP negative somata with branched morphology were manually segmented as astrocytes using Cellpose. For every ROI, median intensity of the SR101 channel was measured in FIJI (Version 2.14.0/1.54f, RRID:SCR_002285). To facilitate comparisons across mice, timepoints, and regions, raw fluorescence measurements were normalized to the median fluorescence intensity of astrocytes within that image.

#### MRI Treatment Study

Male NMRI Nude mice (Charles River) were injected with 500 000 A2058-br cells into the lateral ventricle of the heart. Starting on 5 days post injection, mice were treated daily with intraperitoneal injections of 20 mg kg^-1^ bodyweight MFA (Cayman Chemical, Cat#70550); 10 mg kg^-1^ bodyweight Tonabersat (MedChemExpress, Cat#SB-220453) or vehicle (10% DMSO/PEG400). Littermates were randomly assigned to each treatment group. Treatment was continued until day 28 following intracardiac injection. On day 27 post injection, Gadolinium contrast was administered and a cranial MRI was performed in a 9.4 T, Bruker Topspin 9/20. Mice were sedated with isoflurane and kept under anaesthesia at 0.5–1.5% isoflurane in oxygen. The body temperature was maintained at 37 °C by a heating plate. T1-w and T2-w images were acquired, and tumour volume was quantified through manual segmentation on T1-w images using FIJI (Version 2.14.0/1.54f, RRID:SCR_002285).

#### In Vitro Ca^2+^ Imaging

8 000 tumor cells (DDMel31, A2058-br) or 12 000 tumor cells (NCI-H1915) expressing the genetic calcium indicator GCaMP were seeded on a cell-culture treated glass 96 well plate (Zellkontakt, Cat#5242-20) in their respective media. 48 hours after seeding, medium was exchanged, and a drug or vehicle was added. Plates were transferred to a laser scanning microscope (ZEISS LSM710, LSM900 or LSM980) with an incubator set to 37 degrees Celsius and 5.0% CO_2_. Wells containing drug or vehicle were imaged in random sequence with a 20X/0.8 Plan Apochromat Air objective (ZEISS). GCaMP6s fluorophores were excited using 488nm and recorded using a band-pass emission filter of 490–659 nm with a sampling rate between 0.9 and 2.0 seconds. Laser power was kept below 1% of maximum power to avoid phototoxicity.

The MMBM-MA49 tumoroids were seeded in groups of up to 10 organoids per well into cell culture treated glass 96 well plates and incubated with 1 µM of the chemical calcium indicator Calbryte-520 (ATT Bioquest, Cat#ABD-20650) for 30 minutes prior to imaging and washed once with fresh media. Imaging was performed as mentioned above.

For calcium imaging of A2058-br compared to A2058-par cells, as well as for calcium imaging of A2058-br eGFP-CaProLa, HCT116 and PC3 cell lines, cells were incubated with 1µM of the chemical calcium indicator Calbryte-630 (ATT Bioquest, Cat#ABD-20720) for 30 minutes prior to imaging and washed once with fresh media. Imaging was performed as mentioned above.

Cells were treated with 1µm Senicapoc (Hycultec, Cat#HY-50694) in DMSO (Sigma-Aldrich, Cat#D2438-50ML), 1 µM Tram34 (Hycultec, Cat#HY-13519) in DMSO, 2 mM EGTA (324626-25GM) in nuclease-free water, 1 µM Thapsigargin (biomol, Cat#Cay10522-1) in DMSO, 10 µM BTP2 (Sigma-Aldrich, Cat# Y4895-MG) in DMSO, 200 µM Tonabersat (Hycultec, Cat#HY-15204/CS-0676) in DMSO, 100 µM Carbenoxolone (Hycultec, Cat#HY-B1367) in DMSO and 100 µM Meclofenamate (Cayman Chemical, Cat#70550) in DMSO.

#### Immunofluorescence Staining of MMBM-MA49 tumoroids

Tumoroids were fixed in 4.5% formaldehyde (HistoFix, Carl Roth, Cat#2213.3) for 15 minutes at room temperature and washed three times with PBS (Capricorn Scientific, Cat#PBS-1A). Blocking was performed with 3% BSA (Gerbu, Cat#1063-500g), 0.2% Tween20 (MP Biomedicals, Cat#11TWEEN201-CF) in PBS, followed by overnight incubation at 4°C with mouse anti-Melanoma primary antibody (abcam, Cat#ab732, RRID:AB_305844) at 1:200 in 1% BSA, 0.2% Tween20 in PBS. After three PBS washes, samples were incubated with donkey anti-mouse AF647 secondary antibody (Invitrogen, Cat#A-31571, RRID:AB_162542) at 1:200 in the same buffer for 90 minutes at room temperature. Nuclei were counterstained with DAPI (Sigma-Aldrich, Cat#10236276001).

#### Ca^2+^ Imaging Analysis

<u>Motion Stabilization of In Vivo MPLSM Data</u>: In vivo time-series data was stabilized and motion-corrected with the NoRMCorRE algorithm^117^, using the calcium-independent tdTomato channel. Briefly, outlier images with significant movement were identified by comparing the structural similarity index implemented in scikit-image^122^ (version 0.26.0, RRID:SCR_021142) of consecutive frames. Images with a SSIM < 2 standard deviations of the mean were removed.

<u>Data Preprocessing and Peak Quantification</u>: Analysis of calcium imaging data was performed using the custom-developed pyKrait python package (version 0.3.4), which is freely available at https://pypi.org/project/pykrait/. The following steps were performed within the pyKrait pipeline: A z-projection of the time-series was created and images were normalized using Contrast Limited Adaptive Histogram Equalization (scikit-image, version 0.26.0, RRID:SCR_021142). The resulting 2D projection was segmented fully-automatically for in vitro data sets and semi-automatically in vivo, using custom-trained cellpose models^109,110^ for each cell line (cellpose version 4.6.0, RRID:SCR_021716). The segmentation mask was superimposed onto the time-series and mean fluorescence per ROI per frame was extracted. The resulting traces were detrended using a sinc-filter implemented in PyBoat^118^ (version 1.0.1) and calcium peaks were identified using the find_peaks algorithm implemented in SciPy^124^ (version 1.16.3, RRID:SCR_008058). Peak width was quantified as the full-width at half-maximum (FWHM), calculated using scipy.signal.peak_widths at half the peak prominence height. Rise time was defined as the time elapsed between the 20% and 80% amplitude levels on the ascending part of the peak. Decay time was defined as the time elapsed between the 80% and 20% amplitude levels on the descending part. To analyze overall calcium activity across conditions, calcium peaks were normalized to the number of cells and duration of recording.

Cells in the focal plane whose bounding boxes were within a small pixel distance (3-15 pixels depending on pixel resolution and magnification) of each other were considered adjacent. The neighbor degree of a pair of cells was defined as the shortest path between the two cells and calculated using the floyd warshall algorithm implemented in SciPy^124^ (version 1.16.3, RRID:SCR_008058). For in vivo recordings, only metastases with more than 10 individual cells in the field of view that also showed calcium oscillations were considered in order to ensure a sufficient spatial distribution of activity. Peaks that occurred within 10 frames of each other were considered coinciding, and the total number of such events was counted. As null control, the shortest path matrix was randomly permuted, and the same analysis performed 1000 times. Synchronous Activity was defined as the z-score of experimental data compared to the null control of each data point. To identify pacemaker-like, periodically active cells, the standard deviation of inter-peak intervals was calculated for each cell. Only cells with at least four detected peaks (yielding a minimum of three intervals) were considered. Cells were classified as periodic if their inter-peak interval standard deviation fell below 15 seconds, as described by Hausmann et al.^39^

#### In Silico Modeling of Calcium Oscillations

A kinetic model of the IP_3_ receptor, the Ca^2+^ concentration in the cytoplasm and in the endoplasmic reticulum (ER) as well as, the proportion of IP_3_ receptors that are not inactivated by Ca^2+^ was established by Sneyd et al.^131^ The fluxes J_release_ of Ca from the ER to the cytoplasm and its counterpart J_serca_, the flux of Ca from the cytoplasm into the ER as well as the fluxes of Ca^2+^ out of (J_pm_) and into (J_in_) the cells are used to define a ordinary differential equation (ODE) based in silico model describing the time dynamic of the four above-mentioned quantities. Definitions of the fluxes and the ODEs including their parameters can be found in the Supplementary Material. This model was fitted to a representative region of the calcium dynamic of an example cell. The experimental data, fit region and evaluation of the model with the fitted parameters is shown in Figure 4B. Simultaneous plotting of IP_3_ and Ca^2+^ concentration over time (Figure 4A) shows oscillatory behavior of both quantities, which is qualitatively captured by the model. Inhibition was simulated by multiplying a step function from one to 0.05 with a switch at t=1000 to J_in_ leading to a near instant depletion of cytoplasmic Ca^2+^ concentration (Figure 4C).

#### EdU-Calcium Correlative Imaging

Tumor cells were seeded as described above. After 44 hours, 1µM of EdU (Baseclick GmBH, Cat#BCK-EdUPro488FC50) dissolved in DMSO was added. Calcium Imaging was performed as described previously, and after 4 hours of co-incubation with EdU, cells were fixed with 4.5% formaldehyde (HistoFix, Carl Roth, Cat#2213.3) for 15 minutes at room temperature before washing three times with PBS (Capricorn Scientific, Cat#PBS-1A). EdU staining was performed according to the manufacturer’s manual. The field of view of the calcium imaging was found using reference points and imaging coordinates of the microscope. A zstack of EdU, DAPI and the endogenous fluorescence of tumor cells was acquired. Correlation of cells in EdU staining and calcium imaging datasets were performed manually. EdU positivity was defined by thresholding of the stack histogram.

#### Patch-Clamp Dye Transfer

Whole-cell patch clamp recordings were performed on cultured melanoma cells (DDmel and A2058) plated on glass coverslips (Nunc Thermanox, Cat#174950). Coverslips were transferred to a recording chamber mounted on a fixed-stage upright microscope (BX51WI, Olympus, Hamburg, Germany) and mechanically stabilized using a custom titanium ring to prevent movement during recordings. Cells were continuously perfused with carbonated artificial cerebrospinal fluid (aCSF) maintained at 35 °C. The aCSF contained (in mM): 160 NaCl, 4.5 KCl, 2 CaCl₂, 1 MgCl₂, and 10 HEPES, adjusted to pH 7.4 and an osmolarity of 300 mOsm. Melanoma cells were visualized using differential interference contrast (DIC) optics and epifluorescence illumination. Patch recordings were obtained using a 20× water-immersion objective (XLUMPlan FI, Olympus), and images were acquired with a Hamamatsu EM-CCD camera (C9100–23B) controlled by VisiView software (Visitron Systems GmbH, Puchheim, Germany). For visualization of intracellular dye distribution, fluorescence images were continuously acquired in the GFP channel throughout the recording. Patch electrodes were pulled from borosilicate glass capillaries (1.5 mm outer diameter) and had a resistance of 3–5 MΩ when filled with internal solution. The internal solution contained (in mM): 145 K⁺ aspartate, 2 MgCl₂, 10 HEPES, 10 K₂EGTA, and 8.5 CaCl₂, resulting in a calculated free Ca²⁺ concentration of 1 µM. The pH was adjusted to 7.2 and osmolarity to 290 mOsm. Alexa Fluor 488–conjugated biocytin was included in the internal solution at a concentration of 250 µg/mL to enable visualization of intracellular dye spread. After establishment of the whole-cell configuration, cells were monitored for membrane integrity and dye loading. Fluorescence imaging was performed using a LED-based illumination system (CoolLED pE-4000) with an EGFP 470/40 ET bandpass excitation filter to excite Alexa Fluor 488. Fluorescence images were acquired before, during, and after patching to confirm successful intracellular filling and to assess potential dye diffusion into neighboring cells. Imaging parameters were optimized to maximize temporal resolution while minimizing photobleaching.

#### In Vitro Viability Assay

To assess *in vitro* viability upon gap junction treatment, 50 000 tumor cells per well (A2058, DDmel31, NCI-H-1915) were seeded into an opaque 24 well glass bottom plate (Zell-kontakt, Cat#5241-20). 100µM Meclofenamate or vehicle was added immediately after cell seeding. After 48h of incubation, the CellTiter-Glo (Promega GmbH, Cat#G755) was performed according to the manufacturer’s instructions and were measured using a plate reader (Promega GmbH, GloMax® Explorer Multimode Microplate Reader).

#### Cell Cycle Profiling by EdU Incorporation and DNA Content Staining

50 000 cells were seeded per well in 6-well plates (Faust Lab, Cat#TPP92006) and cultured for 48 h. 36 h after seeding (12 h before fixation), cells were treated with 100 μM meclofenamic acid (Cayman Chemical, Cat#70550), 1µM Senicapoc (Hycultec, Cat#HY-50694), 1µM Tram34 (Hycultec, Cat#HY-13519) or vehicle control for the respective experiments. 44 hours after seeding (4 h before fixation), 1 μM EdU (Baseclick GmbH, Cat#BCK-EdUPro488FC50) was added.

48 h post seeding culture media was collected, cells were washed once with PBS (Capricorn Scientific, Cat#PBS-1A) and dissociated with trypsin (Gibco, Cat#25200-56). Cell suspensions were transferred to 1.5-ml tubes and centrifuged at 500 × g for 5 min. Fixation and EdU detection were performed according to the manufacturer’s Click-iT protocol. Pellets were resuspended in 200 μl PBS, and FxCycle Violet (Invitrogen, F10347) was added at a 1:100 dilution.

Samples were incubated for 20 min at room temperature protected from light, transferred to FACS tubes (Neolab, Cat#352235), and analyzed on a BD FACSCanto II operated at low flow rate. DNA content was detected in the V450/50 (linear) channel (FxCycle), and EdU incorporation in the B530/30 (log) channel (AF488-EdU). Single-stained controls were used for compensation.

Single cells were identified using FSC-A versus SSC-A, and doublets were excluded using SSC-A versus SSC-W. FxCycle intensity was used to distinguish sub-G1, G1, and G2/M populations, and EdU incorporation defined S-phase cells. Data was analyzed using FlowJo (RRID:SCR_008520).

#### Generation of CaProLa_5_^high^ and -^low^ sublines

A2058-br CaProLa_5_ cells were seeded in T175 flasks. After 48 h, culture medium was replaced with fresh medium containing 250 nM CPY-CA dye and incubated for 30 min at 37°C. Dye incorporation was terminated by exchanging the medium for 1 μM recombinant HaloTag enzyme for 5 min. Tumor cells were FACS-isolated based on their CPY-CA/GFP ratio as described above, and re-cultured as CaProLa_5_^high^ and -^low^ subpopulations, respectively. This process was repeated multiple times to enrich the respective subpopulations.

#### CaProLa_5_-Based Profiling, Flow Cytometry and RNA isolation for bulk RNAseq

A2058 CaProLa5 and DDMel31 CaProLa5 cells were seeded at densities of 1–10 million cells in T175 flasks. After 48 h, culture medium was replaced with fresh medium containing 250 nM CPY-CA dye and incubated for 30 min at 37°C. Dye incorporation was terminated by exchanging the medium for 1 μM recombinant HaloTag enzyme for 5 min. Cells were washed twice with PBS, dissociated with trypsin, neutralized with medium, and transferred to 15 ml tubes.

To preserve RNA integrity in fixed cells, the MARIS protocol was adapted^132^. Pellets were centrifuged at 500 × g for 5 min and resuspended in 200 μl 4.5% Formaldehyde supplemented with 80 U RNasin Plus (Promega, Cat#N2615) for 15 min at room temperature. Cells were washed once with 200 μl PBS containing 80 U RNasin Plus, centrifuged at 3000 × g for 3 min, and resuspended in 800 μl PBS supplemented with 640 U RNasin Plus for flow cytometry.

Flow cytometric analysis and cell sorting were performed on a BD FACSAria or BD FACSAria Fusion equipped with a 70-μm nozzle. Single cells were identified using FSC-A, SSC-A, and FSC-W parameters. GFP-positive cells were detected using the BL-530/30 channel, and CaProLa-dependent CPY-CA fluorescence was recorded using the R-670/30 channel. CaProLa-ON and CaProLa-OFF populations served as positive and negative controls to define background fluorescence and signal saturation. For sorting, cells were collected into tubes preloaded with 100 μl PBS containing 160 U RNasin Plus.

Cells sorted from CaProLa-based profiling were centrifuged at 3000 × g for 3 min at 4°C. RNA was isolated using the RecoverAll Total Nucleic Acid Isolation Kit (ThermoScientific, Cat#AM1975) beginning at the protease digestion step. The manufacturer’s protocol was modified by omitting the deparaffinization step and extending protease digestion to 1 h at 50°C. RNA was eluted in 50 μl nuclease-free water (Ambion, Cat#AM9932). RNA concentration was measured with a NanoDrop 1000, and RNA integrity was validated using the 4150 Tapestation System (Agilent, Santa Clara, California, USA, #G2992AA). Library preparation and RNA sequencing were carried out by the GPCF at the DKFZ on a NovaSeq6000 device (Illumina).

#### RNA-Sequencing of Meclofenamate-Treated MBrM Cell Lines

For successive calcium imaging and RNA-Sequencing, 25k A2058-br, 30k DDMel31 or NCI-H1915 cells were seeded in their respective media in a 24 imaging well plate (Zellkontakt, Cat#5242-20). Medium was changed after 36h and 100µM Meclofenamate (Cayman Chemical, Cat#70550) or vehicle was added to the wells. Calcium imaging was performed as described above immediately after the addition of drugs.

48h after seeding (corresponding to 12h of MFA treatment), cells were washed with ice-cold PBS and lysed using the Zymo RNA lysis buffer (Zymo Research Corporation, Cat#R1060-1-50). RNA was isolated using the Zymo Quick RNA 96 kit (Zymo Research Corporation, Cat#R1052) according to the manufacturer’s instructions. On-column DNA digestion was performed with the RNase-Free DNase Set (Qiagen, Cat#79254).

RNA integrity was validated using the 4150 Tapestation System (Agilent, Santa Clara, California, USA, #G2992AA). Library preparation and RNA sequencing were carried out by the GPCF at the DKFZ on a NovaSeq6000 device (Illumina).

#### Analysis of bulk-RNA sequencing data

Reads were aligned and mapped using the One-Touch-Pipeline of the ODCF at the DKFZ^133^. The resulting count matrix was processed using DESeq2^112^ (version 1.44.0, RRID:SCR_015687) to derive differentially expressed genes.

KCNN4 expression between brain metastasis cell lines and glioblastoma cell lines was compared by generating a combined DESeq2 model and analyzing VST-normalized counts. The glioblastoma RNA Sequencing data was obtained from EGAS50000000477 and previously published by Azorin et al.^33^

To generate the MBrM Calcium Signature Score, pairwise differential expression analyses were performed for each combination of CaProLa condition (high, medium, low) and cell line (A2058, DDMel31) using DESeq2. Genes were considered differentially expressed at an adjusted p-value < 0.05 and absolute log2 fold change > 0.5. To derive a consensus signature, genes were required to be significantly differentially expressed in at least one pairwise comparison in each cell line independently, and to show concordant directional regulation (exclusively up- or downregulated) across all comparisons in which they were significant. Batch effects attributable to cell line were removed from VST-normalized counts using the removeBatchEffect function implemented in limma^116^ (version 3.60.6, RRID:SCR_010943), and z-scores were computed per gene within each cell line separately prior to visualization with ComplexHeatmap^111^ (version 2.20.0, RRID:SCR_017270).

Gene set enrichment analysis (GSEA) was performed on a combined DESeq2 model taking into account both cell line and condition using the gsePathway function implemented in ReactomePA^120^ (version 1.48.0, RRID:SCR_019316) Pairwise pathway similarity was computed via enrichplot’s pairwise_termsim (version 1.24.4, RRID:SCR_026996). Significant pathways were represented as a network graph in which nodes correspond to pathways and edge weights reflect term similarity. Cluster detection was performed using the Leiden algorithm via igraph^115^ (version 2.1.4, RRID:SCR_019225) and cluster consensus names were derived manually.

Signature scoring was performed using the singscore package^126^ (version 1.24.0). Counts corrected for cell-line batch effects (VST-normalized) were rank-transformed with rankGenes, and simpleScore was applied to compute single-sample enrichment scores. The MBrM Calcium Signature is described above, the GBM Calcium Signature was derived by Hai, Hoffmann et al.^57^ and the IEG, DPRG and SRG signatures were developed by Tullai et al.^52^. Cell cycle phase enrichment based on the Whitfield et al.^56^ gene sets was assessed by GSEA using the clusterProfiler (version 4.12.6, RRID:SCR_016884) GSEA function.

#### Analysis of MBrM Calcium Signature in brain-metastatic cell lines

Raw counts of brain-tropic and parental cell lines were obtained from GSE83132 and are published in Boire et al.^77^. A DESeq2 model incorporating cell line and condition (parental vs. brain metastatic) was fitted, and VST-normalized expression values were obtained. Gene annotation was performed via biomaRt^108^ (version 2.60.1, RRID:SCR_019214) querying Ensembl using Entrez gene identifiers. Signature scoring on VST-normalized counts was was performed as described above.

#### Analysis of MBrM Calcium Signature in a patient scRNA-Seq Dataset

scRNA-Sequencing data of patient melanoma brain metastases and extracranial metastases samples were obtained from GSE200218 and are published in Biermann et al^26^. Analysis of the MBrM Calcium Signature was restricted to tumor cells with more than 100 detected genes. Counts were normalized, log1p-transformed, and scaled. Signature scoring was performed at the single-cell level using scanpy’s score_genes function^134^ (version 1.1.15, RRID:SCR_018139). Up- and downregulated signature components were scored separately and z-score standardized, and a composite score was derived by subtracting the scaled downregulated score from the scaled upregulated score. Patient-level mean scores were computed by averaging single-cell scores across all tumor cells per patient.

#### Ion Channel Subgroup Analysis of Healthy Brain and MBrM Tissue

RNA-sequencing data of a cohort of healthy brain tissue and MBrM tissue was previously published^60,61^. Counts per million were accessed, ion channels were subset and a differential gene expression analysis with limma^116^ (version 3.60.6, RRID:SCR_010943) was performed.

#### Survival Analysis in a Melanoma Patient Dataset

Gene expression data were obtained from the Gene Expression Omnibus (GEO) database (accession GSE65904^78^) using the GEOquery^114^ (version 2.72.0, RRID:SCR_000146) package. Probe-level expression values were mapped to Ensembl gene identifiers using the illuminaHumanv4.db annotation package. Expression data were log_2_(x + 1) transformed, followed by row-wise z-score normalization across samples. Clinical metadata were extracted from the GEO phenotype data and curated to obtain disease-specific survival time (days), vital status (death = 1, alive = 0), age, sex, and tumor stage. Samples with missing survival, age, or sex data were excluded. Patients were stratified into high and low GJC1 expression groups based on a z-score threshold of 0. Kaplan–Meier survival curves were estimated using the survival package (version 3.8-3, RRID:SCR_026244) and visualized with survminer (version 0.5.0, RRID:SCR_021094). Between-group differences in disease-specific survival were assessed using the log-rank test.

#### 3D Visualization

For 3D visualization of metastases, raw image files were loaded into Aivia (Leica, version 10.5.1) and a pixel probability map for blood vessels and tumors was generated using a custom-trained algorithm. The pixel-probability map was thresholded and smoothed with Fiji (Version 2.14.0/1.54f, RRID:SCR_002285) using a median filter with a pixel-size of 0.5 and converted into STL with a custom python script using scikit-image (scikit-image, version 0.26.0, RRID:SCR_021142) and numpy-stl (version 3.2.0). STL files were imported into blender (version 4.4, RRID:SCR_008606), post-processed and rendered.

### Quantification and statistical analysis

Statistical analyses were performed using GraphPad Prism (version 10.2.2, RRID:SCR_002798), R (version 4.4.1 RRID:SCR_001905) or Python (version 3.11.13, RRID:SCR_008394) using Seaborn^125^ (version 0.13.2, RRID:SCR_018132), SciPy^124^ (version 1.16.3, RRID:SCR_008058), scikit-posthoc^123^ (version 0.11.4, RRID:SCR_021363) and Statsmodels^135^ (version 0.14.6, RRID:SCR_016074). P-values were visualized using Statannotations (version 0.7.2, RRID:SCR_026623). Figures were assembled in Adobe Illustrator 2026 (RRID:SCR_010279).

Normality was assessed with the Shapiro-Wilk test. The exact statistical test, corrections for multiple testing and experimental numbers are indicated in the corresponding figure legends. No statistical methods were used to predetermine sample size. Differences were considered statistically significant at p < 0.05.

---

**Supplementary Figure 1.**
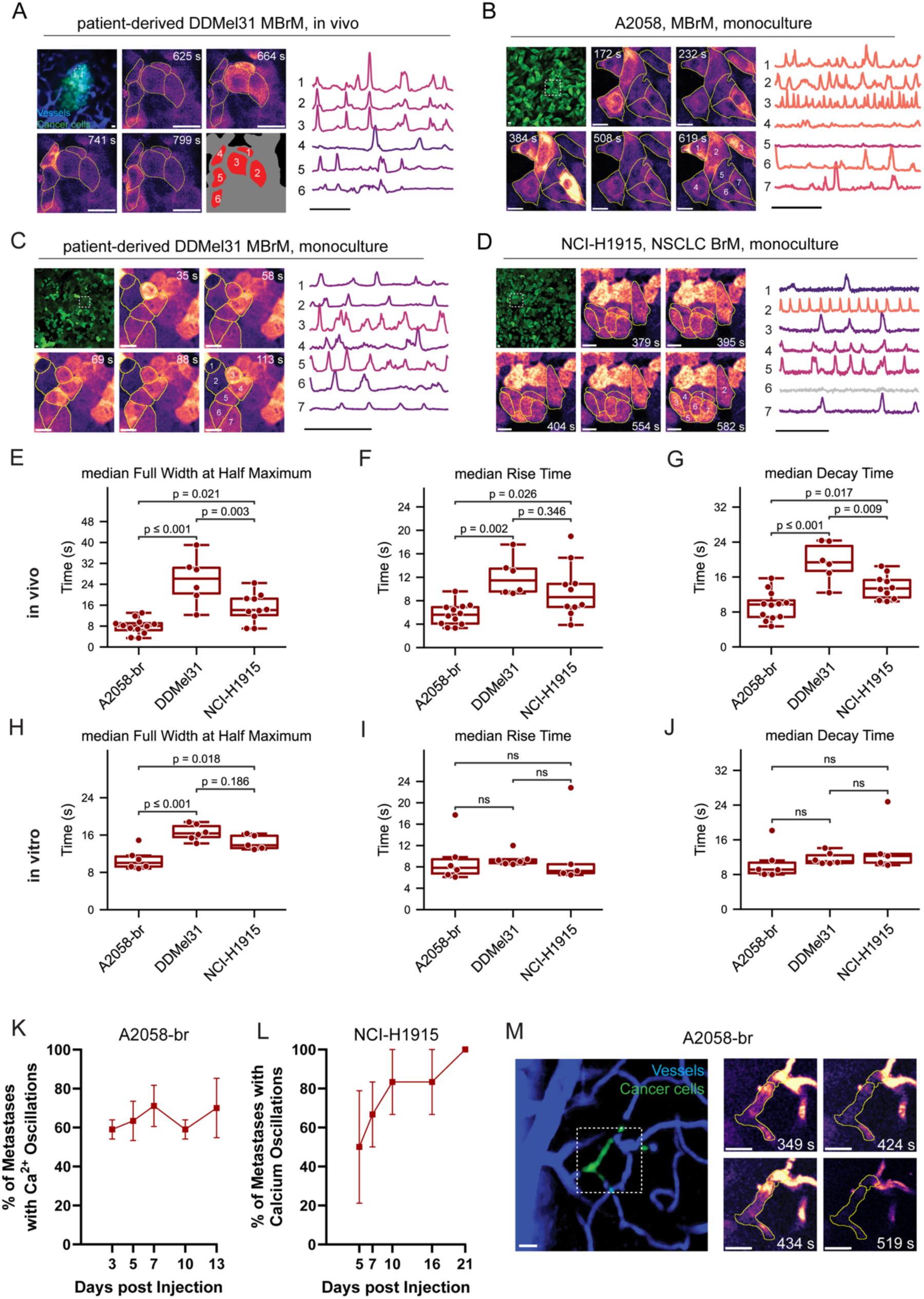
Characterization of Ca^2+^ transients across BrM models. Relates to Figure 1. (A) In vivo Ca^2+^ imaging of the patient-derived DDMel31melanoma brain metastasis after orthotopic engraftment. Top Left: Overview of ROI, Dashed white square indicates zoom-in. Middle: Filmstrip showing GCaMP6s fluorescence intensity over time for the indicated region. Scale bar = 20 µm. Right: Representative Ca^2+^ traces of selected tumor cells over time. Scale bar = 5 min. (B) In vitro monoculture Ca^2+^ imaging of A2058-br brain metastasis cell line. Top Left: Overview of ROI, Dashed white square indicates zoom-in. Middle: Filmstrip showing GCaMP6s fluorescence intensity over time for the indicated region. Scale bar = 20 µm. Right: Representative Ca^2+^ traces of selected tumor cells over time. Scale bar = 5 min. (C) In vitro monoculture Ca^2+^ imaging of DDMel31 brain metastasis cell line. Top Left: Overview of ROI, Dashed white square indicates zoom-in. Middle: Filmstrip showing GCaMP6s fluorescence intensity over time for the indicated region. Scale bar = 20 µm. Right: Representative Ca^2+^ traces of selected tumor cells over time. Scale bar = 5 min. (D) In Vitro Ca^2+^ imaging of NCI-H1915 brain metastasis model. Top Left: Filmstrip of in vitro timelapse recording of the human NSCLC brain metastasis derived NCI-H1915 cell line expressing GcAMP6s. Scalebar = 20µm. Right: Ca^2+^ traces of selected cells over time. Scalebar = 5min. (E) Median Ca2+ peak width (full-width at half-maximum) across models in in-vivo recordings. One-way ANOVA p < 0.001, post-hoc Tukey HSD p-values indicated in figures. n=12/6/10 recordings from n=3/3/3 mice for A2058/DDMel31/NCI-H1915 respectively. (F) Median 20-80% peak rise time across models in in-vivo recordings. One-way ANOVA p = 0.002, post-hoc Tukey HSD p-values indicated in figures. n=12/6/10 recordings from n=3/3/3 mice for A2058/DDMel31/NCI-H1915 respectively. (G) Median 80-20% peak decay time across models in in-vivo recordings. One-way ANOVA p < 0.001, post-hoc Tukey HSD p-values indicated in figures. n=12/6/10 recordings from n=3/3/3 mice for A2058/DDMel31/NCI-H1915 respectively. (H) Quantification of median Ca2+ peak width (full-width at half-maximum) across models in in-vitro recordings. One-way ANOVA p < 0.001 with post-hoc Tukey HSD indicated in figures. n=6/6/5 independent recordings. (I) Quantification of median 20-80% peak rise time across models in in-vitro recordings. Kruskal-Wallis p = 0.318, n = 6/6/5 independent recordings. (J) Quantification of median 80-20% peak decay time across models in in-vitro recordings. Kruskal-Wallis p = 0.190, n = 6/6/5 independent recordings. (K) Percentage of metastases displaying Ca^2+^ oscillations over time after heart injection in the A2058-br model. n=3 mice. Mean and SEM shown. (L) Percentage of metastases displaying Ca^2+^ oscillations over time after heart injection in the NCI-H1915 model. n=3 mice. Mean and SEM shown. (M) Left: A2058-br cancer cell (green) arrested in the brain vasculature (blue). Scale bar = 20 µm. Right: Film-strip of the Ca^2+^ recording. White outline corresponds to the tdTomato tumour signal.

**Supplementary Figure 2.**
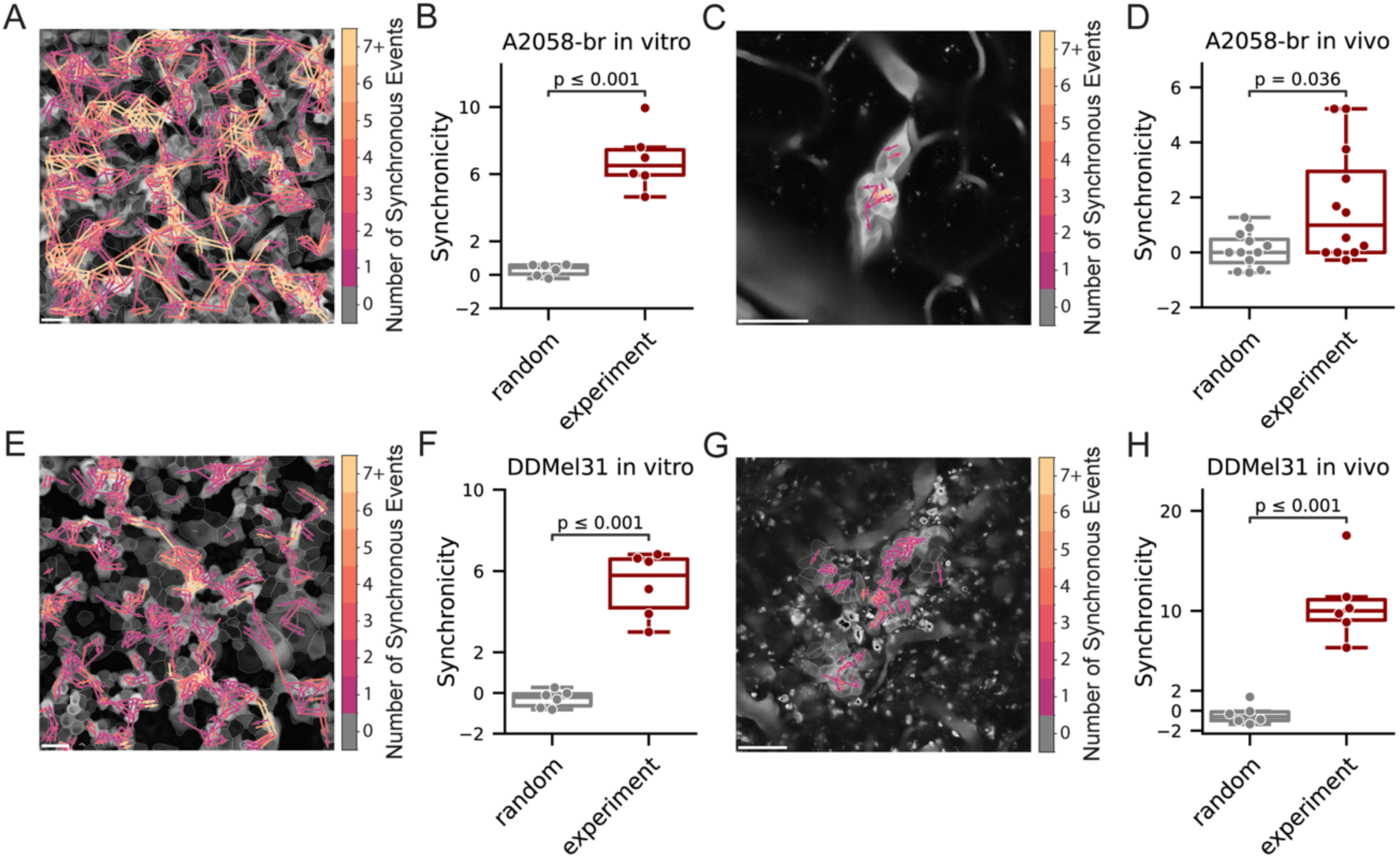
Ca^2+^ Signaling Networks in melanoma BrM models. Relates to Figure 2. (A) Network map of Ca^2+^ activity synchronicity in A2058-br cells in vitro. Connections between neighboring cells are color-coded according to the number of synchronous Ca²⁺ events. Scale bar = 50 µm. (B) Synchronicity of Ca^2+^ activity in A2058-br cells in vitro. Two-sided Welch’s t-test, n = 6 independent recordings. (C) Network map of Ca^2+^ activity synchronicity in A2058-br cells in vivo. Scale bar = 50 µm. (D) Synchronicity of Ca^2+^ activity in A2058-br BrM in vivo. Two-sided Mann-Whitney U test, n=12 recordings from 3 mice. (E) Network map of Ca^2+^ activity synchronicity in DDMel31 cells in vitro. Scale bar = 50 µm. (F) Synchronicity of Ca^2+^ activity in DDMel31 cells in vitro. Two-sided Welch’s test, n=6 independent recordings. (G) Network map of Ca^2+^ activity synchronicity in DDMel31 cells in vivo. Scale bar = 50 µm. (H) Synchronicity of Ca^2+^ activity in DDMel31 BrM in vivo. Two-sided Welch’s t-test, n=6 recordings from 3 mice.

**Supplementary Figure 3.**
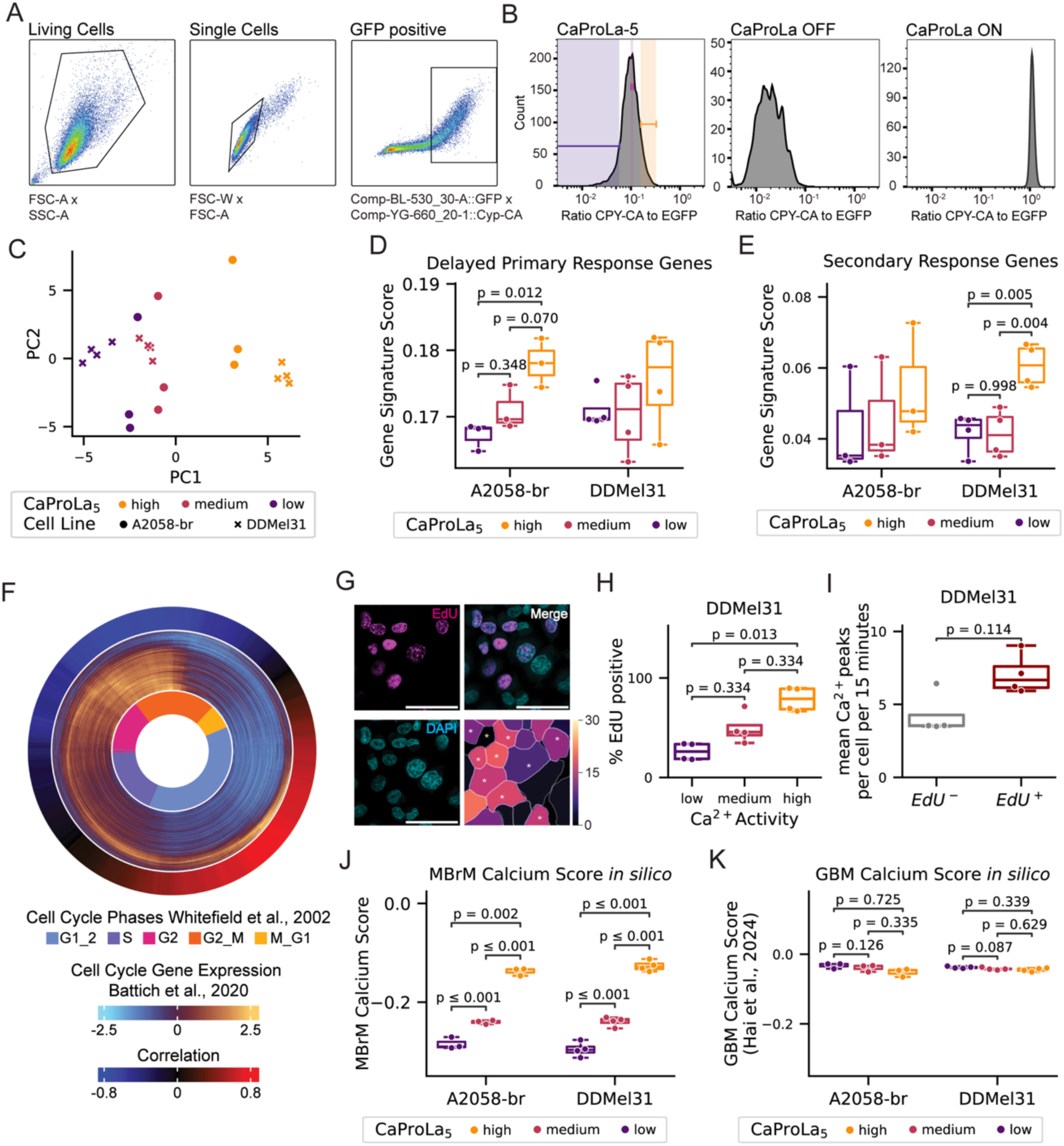
Transcriptomic changes with Ca^2+^ transients across BrM models. Relates to Figure 3. (A) Gating strategy for CaProLa-based sorting. Living cells and singlets were sequentially gated based on FSC/SSC parameters, followed by selection of GFP-positive cells. (B) Representative histograms of CaProLa signal intensity. Distributions of the CPY-CA/GFP ratio are shown for CaProLa-5, CaProLa-OFF, and CaProLa-ON cell lines. Gates for CaProLa-based sorting are indicated in the left panel. (C) Principal Component Analysis of the top 500 variable genes, according to cell line and CaProLa status. n=4 samples per CaProLa status for DDMel31, n=3 samples per CaProLa status for A2058-br. (D) Boxplot of Delayed Primary Response Genes signature score across CaProLa status and models. Two-way ANOVA for cell line (p = 0.899) and CaProLa status (p = 0.031). One-way ANOVA for CaProLa status in A2058-br (p = 0.014) and DDMel31 (p = 0.416). Post-hoc pairwise Tukey’s HSD p-values are shown in the figure. n=3/4 samples for A2058-br and DDMel31, respectively. (E) Boxplot of Secondary Response Genes signature score across CaProLa status and models. Two-way ANOVA for cell line (p = 0.936) and CaProLa status (p = 0.030). One-way ANOVA for CaProLa status in A2058-br (p = 0.675) and DDMel31 (p = 0.002). Post-hoc pairwise Tukey’s HSD p-values are shown in the figure. n=3/4 samples for A2058-br and DDMel31, respectively. (F) Circular heatmap showing correlation coefficients of CaProLa^high^ gene expression fold changes (outer ring) with gene expression fold changes across the cell cycle as reported in Battich et al., 2020 (middle ring) and assigned cell cycle phase according to Whitefield et al., 2002 (inner ring). See methods for details. (G) Correlation of Ca^2+^ Imaging and EdU staining in DDMel31. EdU (magenta, upper left) and DAPI (cyan, bottom left) staining of the same region (middle). Right: Heatmap of Ca^2+^ peaks in 15 minutes of recording. EdU-positive nuclei are annotated with (*). Scalebar = 50µm (H) Percentage of DDMel31 EdU positive cells based on their Ca^2+^ activity status. DDMel31 cells were split into Ca^2+-low^ (0-10th percentile of Ca2+ activity), Ca^2+-medium^ (45-55th percentile) and Ca^2+-high^ (90-100th percentile). Friedman test p = 0.018. Post-hoc Nemenyi p-values are shown in the figure. n=4 independent recordings. (I) Mean Ca^2+^ activity of cells depending on EdU status. Two-sided Man Whitney U test, n=4 independent recordings. (J) Boxplot of MBrM Ca^2+^ Signature score in melanoma brain metastases models. Two-way ANOVA for cell line (p = 0.960) and CaProLa status (p < 0.001). One-way ANOVA for CaProLa status in A2058-br (p < 0.001) and DDMel31 (p < 0.001). Posthoc Tukey’s-HSD p-values are shown in the figure. n=4/3 respectively. (K) GBM Ca^2+^ Signature does not correspond to MBrM CaProLa status, highlighting entity-specific mechanism regulation of Ca^2+^ activity. Two-way ANOVA for cell line (p = 0.949) and CaProLa status (p = 0.020). One-way ANOVA for CaProLa status in A2058-br (p = 0.137) and DDMel31 (p = 0.010). Posthoc Tukey’s-HSD p-values are shown in the figure. n=4/3 respectively.

**Supplementary Figure 4.**
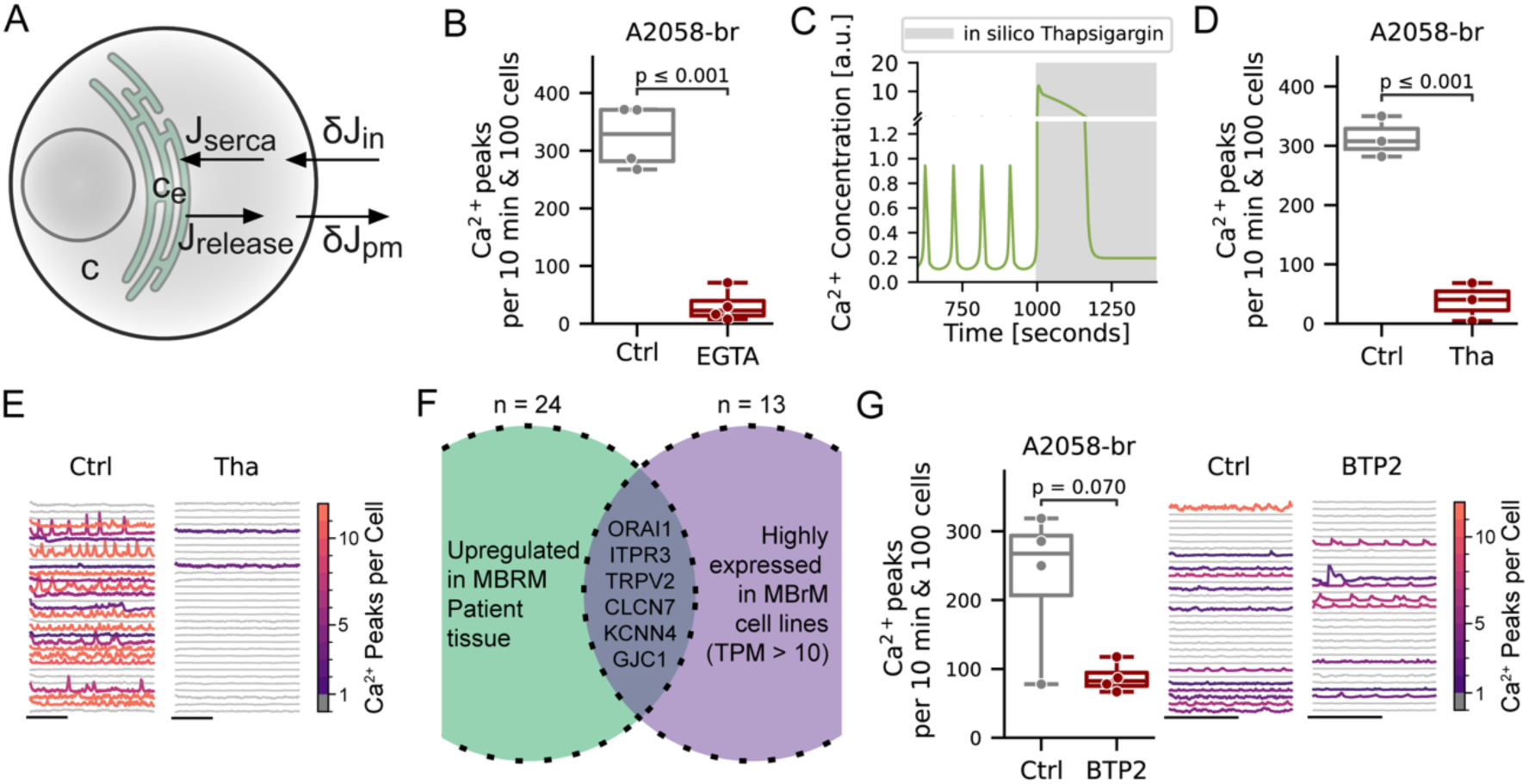
Pathways governing Ca^2+^ release. Relates to Figure 4. (A) Schematic of the mathematical model of cellular Ca^2+^ homeostasis. Four fluxes (J_in_, J_pm_, J_serca_, J_release_) are modelled as depending on intracellular Ca^2+^ and IP_3_ concentration. See methods for details. (B) Chelation of extracellular Ca^2+^ by 2mM EGTA inhibits Ca^2+^ activity in A2058-br melanoma. Two-sided Welch’s t-test, n=4 independent recordings. (C) *In silico* simulation of Ca^2+^ dynamics after extracellular Ca^2+^ chelation with Thapsigargin (grey area), showing rapid loss of Ca^2+^ oscillations (D) Depletion of ER Ca^2+^ with 1µM Thapsigargin inhibits Ca^2+^ activity in A2058-br melanoma. Two-sided Welch’s t-test, n=3 independent recordings. (E) Representative Ca^2+^ traces per cell showing loss of Ca^2+^ activity after 1µM Thapsigargin treatment. Scalebar = 5 minutes. (F) Venn Diagram showing overlap of ion channels upregulated in melanoma brain-metastasis tissue in patients compared to healthy brain, and highly expressed (TPM > 10) in melanoma brain metastasis cell lines. (G) Left: Boxplot showing reduced Ca^2+^ activity in A2058-br cells treated with 10µM BTP2 compared to vehicle. Two-sided Welch’s t-test, n=4 independent recordings. Right: Representative Ca^2+^ traces per cell showing suppression of Ca^2+^ oscillations with 10µM BTP2 treatment. Scalebar = 5 minutes.

**Supplementary Figure 5.**
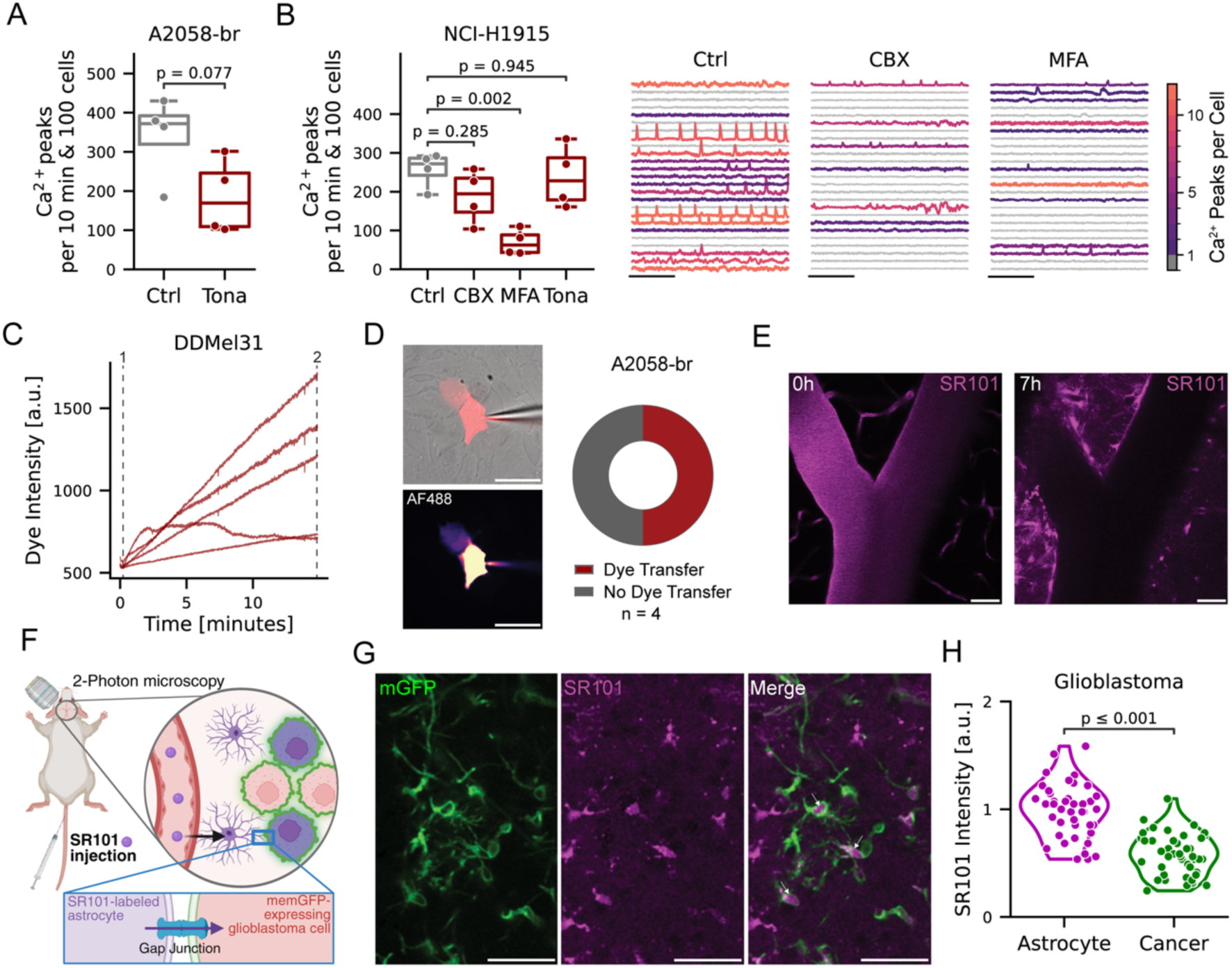
Gap-junction coupled networks in BrM. Relates to Figure 5. (A) Boxplot of Ca^2+^ activity of A2058-br melanoma brain metastases cells after treatment with 200µM tonabersat (Tona) or vehicle. Two-sided Welch’s t-test, n=4. (B) Left: Boxplot of Ca^2+^ activity of NCI-H1915 lung adenocarcinoma cells after treatment with 100 µM carbenoxolone (CBX), 100 µM meclofenamate (MFA) or 200 µM tonabersat (Tona) or vehicle. One-way ANOVA p = 0.004. Post-hoc Dunn’s test with Holm correction p-values are indicated in the figure. Right: Ca^2+^ traces of recordings. Cells are color-coded based on Ca^2+^ activity. Scalebar = 5 minutes. (C) Dye Uptake in neighbouring cells is time-dependent. Line plot of fluorescence intensity of neighbouring cells after patching. (D) Dye transfer between A2058-br melanoma cells. Left: Representative images of a large, membrane impermeable AF488 dye injected into a tumour cell with uptake in a neighbouring cell (arrowhead). Right: Quantification of observed dye transfer in n=4 independent experiments. (E) Left: In Vivo 2-Photon Microscopy of SR101 signal in the vasculature. Right: In Vivo 2-Photon Microscopy of SR101 after uptake by astrocytes. Scalebar = 50µm. (F) Schematic of SR101 astrocyte labelling mechanism in glioblastoma. Created in https://BioRender.com. (G) Representative image of the S24 glioblastoma cell line expressing membrane-targeted GFP (green) and astrocyte staining with SR101 (magenta). Some glioma cells take up the SR101 dye (arrows). (H) Quantification of SR101 fluorescence intensity of individual tumour cells or astrocytes in the S24 glioblastoma model. Tumour cells take up varying concentrations of SR101 dye. n=41 astrocytes and n=48 tumour cells.

**Supplementary Fig. 6.**
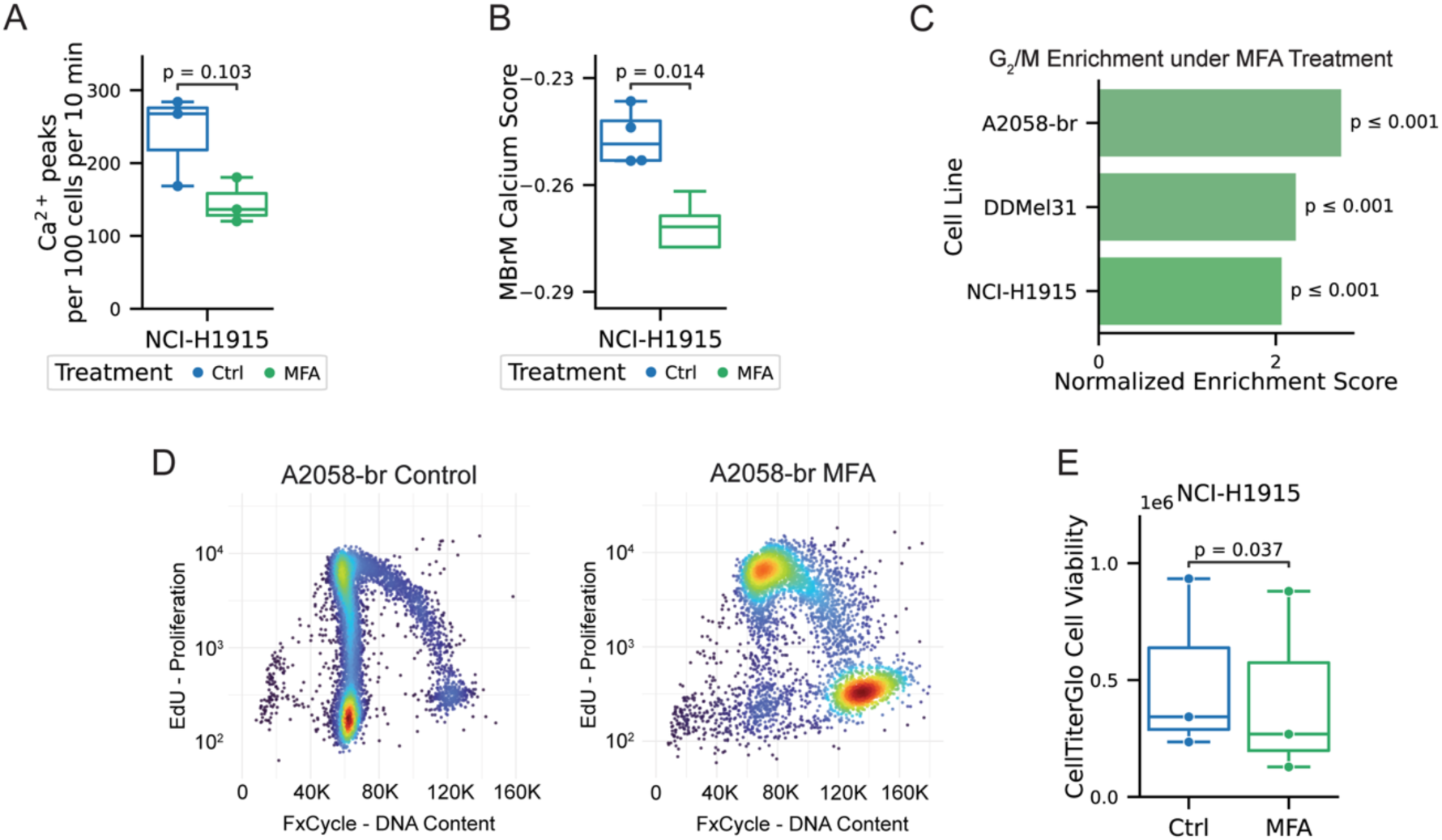
Molecular effects of Ca^2+^ inhibition. Relates to Figure 6. (A) Boxplot of Ca²⁺ activity of NCI-H1915 lung adenocarcinoma samples submitted to RNA-Sequencing treated with control or 100 µM Meclofenamate (MFA). Two-sided Welch’s t-Test, n=3. (B) Boxplot of MBrM Ca^2+^ Signature score in NCI-H1915 lung adenocarcinoma brain metastases models. Two-sided Welch’s t-Test, n=3. (C) Barplot showing gene set enrichment analysis for the G_2_/M gene set per cell line under MFA treatment. (D) Cell Cycle FACS of A2058-br 12 hours after 100 µM MFA treatment, DNA content labelled with FxCycle and proliferation status with EdU. (E) Boxplot of cell viability after 48 hours of vehicle versus MFA treatment in NCI-H1915 cells, measured via CellTiterGlo assay. n = 3 independent biological experiments. Two-sided Welch’s t-test.

**Supplementary Fig. 7.**
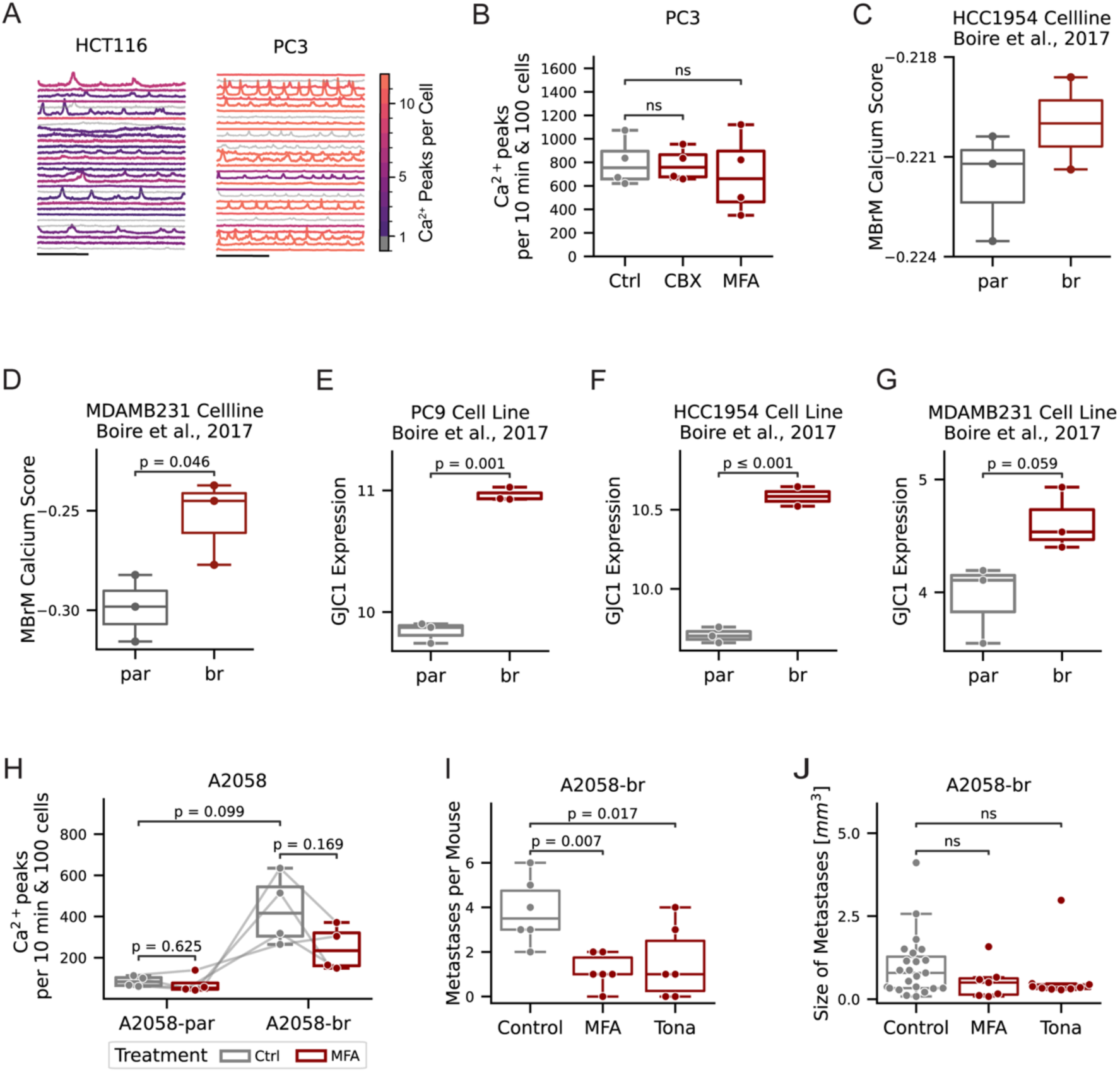
Extended characterisation of Ca²⁺ signalling and gap junction expression across cancer types and metastatic models. Relates to Figure 7. (A) Left: Traces of Ca^2+^ activity in colorectal cancer cell line HCT116. Right: Traces of Ca^2+^ activity in prostate cancer cell line PC3. (B) Gap junction inhibition does not affect Ca^2+^ activity in PC3. One-way ANOVA p = 0.823. (C) Comparison of in-silico MBrM Ca^2+^ signature score of the parental (par) or brain-tropic (br) HCC1954 breast cancer cell line. Raw transcriptomic data obtained from GSE83132. n=3/2. (D) Comparison of in-silico MBrM Ca^2+^ signature score of the parental (par) or brain-tropic (br) MDAMB231 breast cancer cell line. Raw transcriptomic data obtained from GSE83132. n=3, Welch’s t-test. (E) Expression of gap junction gene GJC1 of the parental (par) or brain-tropic (br) PC9 lung cancer cell line in vitro. Raw transcriptomic data obtained from GSE83132. Variance-stabilization transformed (vst) count data shown. Two-sided Welch’s t-test, n=3 (F) Expression of gap junction gene GJC1 of the parental (par) or brain-tropic (br) HCC1954 breast cancer cell line in vitro. Raw transcriptomic data obtained from GSE83132. Variance-stabilization transformed (vst) count data shown. Two-sided Welch’s t-test, n=3/2 (G) Expression of gap junction gene GJC1 of the parental (par) or brain-tropic (br) MDAMB231 breast cancer cell line in vitro. Raw transcriptomic data obtained from GSE83132. Variance-stabilization transformed (vst) count data shown. Two-sided Welch’s t-test, n=3 (H) Ca^2+^ activity in A2058-par and A2058-br cells upon treatment with 100 µM Meclofenamate. Two-sided paired Wilcoxon or t-test, corrected for multiple testing with Benjamini-Hochberg, n=4. (I) Quantification of number of A2058 brain metastases per mouse on day 28 post heart injection measured by MRI. One-way ANOVA p = 0.007 with post-hoc Dunnett’s test p-values indicated in figure. n=6 mice per group. (J) Quantification of the individual size of A2058 brain metastases on day 28 post heart injection measured by MRI. Kruskal-Wallis p-value = 0.259. n=23/7/9 metastases per group.

