## Supplementary Methods for "Collective cancer cell calcium activity drives brain metastasis"

### Materials and Methods (Appendix)

In Sneyd et al., the fluxes are defined as

$$
\begin{aligned}
J_{\text{release}} &= k_{\text{flux}}\left(\mu_0 + \frac{\mu_1 p}{k_{\text{mu}} + p}\right) n \left(b + \frac{V_1 C}{k_1 + C}\right)(C_e - C) \\
J_{\text{serca}} &= \frac{V_e C}{k_e + C} \\
J_{\text{pm}} &= \frac{V_p C^2}{k_p^2 + C^2} \\
J_{\text{in}} &= \alpha_1 + \alpha_2 \frac{\nu}{\beta}.
\end{aligned}
$$

The ODEs are:

$$
\begin{aligned}
\frac{\mathrm{d}C}{\mathrm{d}t} &= J_{\text{release}} - J_{\text{serca}} + \delta(J_{\text{in}} - J_{\text{pm}}) \\
\frac{\mathrm{d}C_e}{\mathrm{d}t} &= \gamma(J_{\text{serca}} - J_{\text{release}}) \\
\frac{\mathrm{d}n}{\mathrm{d}t} &= \frac{1}{\tau}\left(\frac{k_2^2}{k_2^2 + C^2} - n\right) \\
\frac{\mathrm{d}p}{\mathrm{d}t} &= \nu \frac{C + (1-\alpha)k_4}{C + k_4} - \beta p.
\end{aligned}
$$

Both, the fluxes and ODEs are taken from Sneyd et al. The model parameters, their biological meaning and used values are shown in table 1.

**In silico inhibition** To simulate inhibition of $J_{\text{serca}}$ and $J_{\text{in}}$, a smooth, near-instant suppression curve $f_{\text{sup}}$ that switches from one to 0.05 at a specified switch time point is multiplied to the respective flux in the evaluation of the ODE model:

$$
f_{\text{sup}}(t) = 0.05 + 0.95 \frac{1 + \tanh(-k(t - t_{\text{switch}}))}{2}, \quad k = 100, \quad t_{\text{switch}} = 1000. \tag{1}
$$

At $t_{\text{switch}} = 1000$, the function switches from one to 0.05 with a steepness controlled by $k = 100$. This construction ensures, that the switch fulfills differentiability requirements necessary for the fitting process.

### References

J. Sneyd, K. Tsaneva-Atanasova, V. Reznikov, Y. Bai, M. J. Sanderson, and D. I. Yule. A method for determining the dependence of calcium oscillations on inositol trisphosphate oscillations. 103(6):1675–1680. ISSN 0027-8424, 1091-6490. doi: 10.1073/pnas.0506135103. URL `https://pnas.org/doi/full/10.1073/pnas.0506135103`.

Table 1: Complete parameter set for model evaluation

| **Symbol** | **Value** | **Unit** | **Biological function** |
|---|---|---|---|
| *Fitted parameters (pars vector)* | | | |
| $V_e$ | 9.13 | $\mu$M/s | Maximum pumping rate into the ER |
| $k_e$ | 0.153 | $\mu$M | $Ca^{2+}$ level for half-max ER pumping |
| $V_p$ | 11.82 | $\mu$M/s | Maximum pumping rate from the cytosol |
| $k_p$ | 0.351 | $\mu$M | $Ca^{2+}$ level for half-max cytosolic pumping |
| $k_\mu$ | 1.87 | $\mu$M | Half-max concentration parameter in $J_{\text{release}}$ |
| $k_{\text{flux}}$ | 0.493 | $\mu$M/s | Maximum total $Ca^{2+}$ flux through $IP_3$Rs |
| $k_1$ | 0.291 | $\mu$M | $Ca^{2+}$ sensitivity constant for $IP_3$R activation in $J_{\text{release}}$ |
| $k_2$ | 3.75 | $\mu$M | $Ca^{2+}$ concentration scale controlling $IP_3$R inactivation |
| $k_4$ | 0.250 | $\mu$M | $Ca^{2+}$ sensitivity constant in $IP_3$ production dynamics |
| $\alpha_1$ | 2.78 | $\mu$M/s | Basal $Ca^{2+}$ influx rate |
| $\alpha_2$ | $7.2 \times 10^{-14}$ | $s^{-1}$ | $IP_3$-dependent scaling of $Ca^{2+}$ influx |
| $\beta$ | 0.0795 | $s^{-1}$ | Constant rate of $Ca^{2+}$ influx into cytosol |
| $\gamma$ | 0.461 | – | Ratio of cytosolic volume to ER volume |
| $\delta$ | 0.0359 | – | Scale factor: PM $Ca^{2+}$ flux / ER flux |
| *Fixed parameters (hard-coded in model)* | | | |
| $\mu_0$ | 0.567 | – | Proportion of $IP_3$R activated without $Ca^{2+}$ |
| $\mu_1$ | 0.433 | – | Proportion of $IP_3$R activated by $IP_3$ |
| $b$ | 0.111 | – | Basal $IP_3$R opening probability |
| $V_1$ | 0.889 | – | $Ca^{2+}$-dependent $IP_3$R activation fraction |
| $\alpha$ | 1 | – | Weighting of $Ca^{2+}$ feedback on $IP_3$ production |
| $\tau$ | 2 | s | Time scale of $IP_3$R inactivation |
| $\nu$ | 0.56 | $s^{-1}$ | $IP_3$ production rate parameter |
| *Initial conditions* | | | |
| $C(0)$ | 0.0324 | $\mu$M | Initial cytosolic $Ca^{2+}$ concentration |
| $C_e(0)$ | 26.47 | $\mu$M | Initial ER $Ca^{2+}$ concentration |
| $n(0)$ | 0.438 | – | Initial fraction of non-inactivated $IP_3$Rs |
| $p(0)$ | 2.655 | $\mu$M | Initial $IP_3$ concentration |
